# cGAMP loading enhances the immunogenicity of VLP vaccines

**DOI:** 10.1101/2020.01.03.893586

**Authors:** Lise Chauveau, Anne Bridgeman, Tiong Kit Tan, Ryan Beveridge, Joe Frost, Isabela Pedroza-Pacheco, Thomas Partridge, Persephone Borrow, Hal Drakesmith, Alain Townsend, Jan Rehwinkel

## Abstract

Cyclic GMP-AMP (cGAMP) is an immunostimulatory second messenger produced by cGAS that activates STING. Soluble cGAMP acts as an adjuvant when administered with antigens. cGAMP is also incorporated into enveloped virus particles during budding. We hypothesised that inclusion of the adjuvant cGAMP within viral vaccine vectors would promote adaptive immunity against vector antigens. We immunised mice with virus-like particles (VLPs) containing the HIV-1 Gag protein and VSV-G. Inclusion of cGAMP within these VLPs augmented splenic VLP-specific CD4 and CD8 T cell responses. It also increased VLP- and VSV-G-specific serum antibody titres and enhanced *in vitro* virus neutralisation. The superior antibody response was accompanied by increased numbers of T follicular helper cells in draining lymph nodes. Vaccination with cGAMP-loaded VLPs containing haemagglutinin induced high titres of influenza A virus neutralising antibodies and conferred protection following subsequent influenza A virus challenge. Together, these results show that incorporating cGAMP into VLPs enhances their immunogenicity, making cGAMP-VLPs an attractive platform for novel vaccination strategies.

**Short summary:** cGAMP is an innate immune signalling molecule that can be transmitted between cells by inclusion in enveloped virions. This study demonstrates enhanced immunogenicity of HIV-derived virus-like particles containing cGAMP. Viral vectors loaded with cGAMP may thus be potent vaccines.

## Introduction

Vaccination is a powerful strategy in the fight against infectious disease, including virus infection. Indeed, vaccination led to the global eradication of smallpox and is higly protective against some viruses including measles virus and yellow fever virus. However, the development of vaccines inducing long-lasting and broadly effective protection has been difficult for other viruses such as human immunodeficiency virus (HIV) and influenza A virus (IAV), highlighting the need for new vaccination strategies (Rappuoli et al., 2011).

Successful vaccines induce potent adaptive immune responses. Prophylactic vaccine-mediated protection against most virus infections is thought to be predominantly due to induction of antiviral antibody responses that prevent or rapidly control subsequent infection. Antibody responses limit virus infection and spread through several mechanisms (Pelegrin et al., 2015). Neutralising antibodies directly bind virus particles and prevent them from infecting cells. Virion-bound antibodies can also trigger complement activation and virolysis. In addition, antibodies can bind to virus-infected cells and target them for lysis or viral clearance by complement or cells mediating cytolytic or viral inhibitory activity such as natural killer cells and macrophages. Following immunisation, antibodies are initially produced by short-lived extrafollicular plasmablasts. To achieve long-term protection, long-lived plasma cells and memory B cells must be generated in secondary lymphoid tissues. This process occurs in specialised structures called germinal centres (GCs) (Cyster and Allen, 2019; Linterman and Hill, 2016). In GCs, a complex interplay between follicular dendritic cells (FDCs), tingible body macrophages, CD4 T follicular helper (Tfh) cells, CD4 T follicular regulatory (Tfr) cells and B cells results in the formation of long-lived plasma cells and memory B cells producing high-affinity antibodies that confer durable protection. Tfh cells are a CD4 T cell subset specialised to provide help to B cells and are essential for GC formation. They increase the magnitude and quality of the humoral response by promoting B cell proliferation, isotype switching and plasma cell differentiation; by mediating selection of high-affinity B cells in GCs; and by supporting the generation of long-lived plasma cells and memory B cells (Crotty, 2019). In contrast, CD4 Tfr cells are involved in limiting GC reactions to prevent autoantibody formation. Therefore, the Tfh/Tfr ratio is important for regulation of GC responses (Sage et al., 2013).

Virus-specific cytotoxic T cell (CTL) responses mediate clearance of infected cells to prevent virus spread and eradicate infection. If sterilising immunity is not conferred by antibodies, CTLs can make a key contribution to prophylactic vaccine efficacy (Hansen et al., 2011), and they are critical for the control of persistent infection with viruses such as hepatitis B virus, hepatitis C virus, cytomegalovirus and HIV (Panagioti et al., 2018). CTLs exert their activity by triggering destruction of infected cells via release of perforins and granzymes; by ligation of death-domain containing receptors and/or secretion of TNFα; and by producing “curative” cytokines such as IFNγ. Both the magnitude, i.e. the number of activated cells, and polyfunctionality of the T cell response, i.e. the capacity to mediate a breadth of effector activities including production of multiple cytokines, are important determinants of CD8 T cell-based vaccine efficacy (Panagioti et al., 2018).

Initiation of virus-specific CD4 and CD8 T cell responses requires presentation of viral antigens to naïve T cells by professional antigen-presenting cells (APCs), principally dendritic cells (DCs). T cells need to receive three signals for activation: T cell receptor (TCR) triggering by contact with peptide-major histocompatibility complexes (MHC) (signal 1); costimulatory signals (signal 2); and inflammatory cytokines (signal 3) (Joffre et al., 2009).

To induce adaptive immune responses, vaccines need to contain not only appropriate antigens but also an adjuvant. Adjuvants exert a breath of effects; for example, they induce the expression of costimulatory molecules and cytokines by DCs (Coffman et al., 2010). There are only a limited number of FDA-approved adjuvants, most of which are based on aluminium salts (Shi et al., 2019). The increasing knowledge in the field of innate immunity, particularly in the mechanisms underlying pathogen recognition by innate immune receptors, provides an opportunity to develop new adjuvants that specifically engage such receptors and trigger a robust response (Temizoz et al., 2018). Adjuvants targeting toll-like receptors or the cytosolic DNA sensing pathway have attracted a lot of attention (Dubensky et al., 2013). In particular, cyclic dinucleotides (CDNs) that activate stimulator of interferon genes (STING, also known as TMEM173, MPYS, ERIS and MITA) and induce a type I interferon (IFN-I) response as well as production of pro-inflammatory cytokines are being developed as adjuvants (Cai et al., 2014). CDNs facilitate both CD8 T cell and antibody responses (Blaauboer et al., 2014; Kuse et al., 2019; Li et al., 2013) and are effective as mucosal adjuvants (Blaauboer et al., 2015; Ebensen et al., 2011). 2’-3’ cyclic GMP-AMP (cGAMP) is of particular interest. It is produced by cGAMP synthase (cGAS) upon DNA sensing in the cell cytoplasm (Ablasser et al., 2013; Diner et al., 2013; Sun et al., 2013). Soluble cGAMP has been employed as an adjuvant in multiple pre-clinical vaccination models and is an anti-tumour agent (Corrales et al., 2015; Demaria et al., 2015; Li et al., 2016; Li et al., 2013; Wang et al., 2017). However, cGAMP levels are likely to diminish quickly in the extracellular milieu, due to diffusion from the site of administration and degradation by phosphodiesterases such as Ectonucleotide Pyrophosphatase/Phosphodiesterase 1 (ENPP1), an enzyme degrading extracellular ATP and cGAMP (Carozza et al., 2019; Li et al., 2014). Indeed, when injected intra-muscularly, the concentration of cGAMP at the inoculation site decreases rapidly, resulting in a sub-optimal adjuvant effect (Wang et al., 2016).

We and others previously showed that cGAMP is packaged into nascent viral particles as they bud from the membrane of an infected cell (Bridgeman et al., 2015; Gentili et al., 2015). Upon virus entry into newly infected cells, cGAMP is released into the cytosol and directly activates STING. Building on this observation, we hypothesised that inclusion of the adjuvant cGAMP in viral vaccine vectors may enhance their immunogenicity by targeting adjuvant and antigen to the same cell and by protecting cGAMP from degradation in the extracellular environment.

Indeed, using HIV-derived viral-like particles (VLPs), we found that the presence of cGAMP within VLPs enhanced adaptive immune responses to VLP antigens. Antigen-specific CD4 and CD8 T cell responses were augmented, as well as neutralising antibody production. The latter was accompanied by an increase in Tfh cells in draining lymph nodes. cGAMP-loaded VLPs containing the IAV haemagglutinin protein induced neutralising antibodies and conferred protection against development of severe disease after challenge with live IAV. These results highlight the utility of cGAMP loading as a strategy to boost the immunogenicity of viral vaccine vectors.

## Results

### cGAMP-loading of HIV-derived VLPs

HIV-derived viral vectors and VLPs are routinely produced in the cell line HEK293T by transfection of plasmids encoding viral components (Milone and O’Doherty, 2018). Here, we generated VLPs by using plasmids expressing the HIV-1 capsid protein Gag fused to GFP (Gag-GFP) and the Vesicular Stomatitis Virus envelope glycoprotein (VSV-G). The resulting VLPs consist of a Gag-GFP core and a lipid membrane derived from the producer cell that is spiked with VSV-G proteins. Of note, these VLPs do not contain viral nucleic acid and can therefore not replicate in the host (Deml et al., 2005). Additional over-expression of cGAS in the VLP producer cells results in its activation, presumably by the transfected plasmid DNA, and in the presence of cGAMP in the cytosol. It is noteworthy that HEK293T cells do not express STING (Burdette et al., 2011); therefore, cGAS-overexpressing VLP producer cells do not respond to the presence of cGAMP. cGAMP is then packaged into the nascent viral particles, which are released as cGAMP-loaded VLPs (hereafter cGAMP-VLPs; Fig 1A). As a control, we produced VLPs that do not contain cGAMP (Empty-VLPs) by using a catalytically inactive version of cGAS.

**Fig 1:**
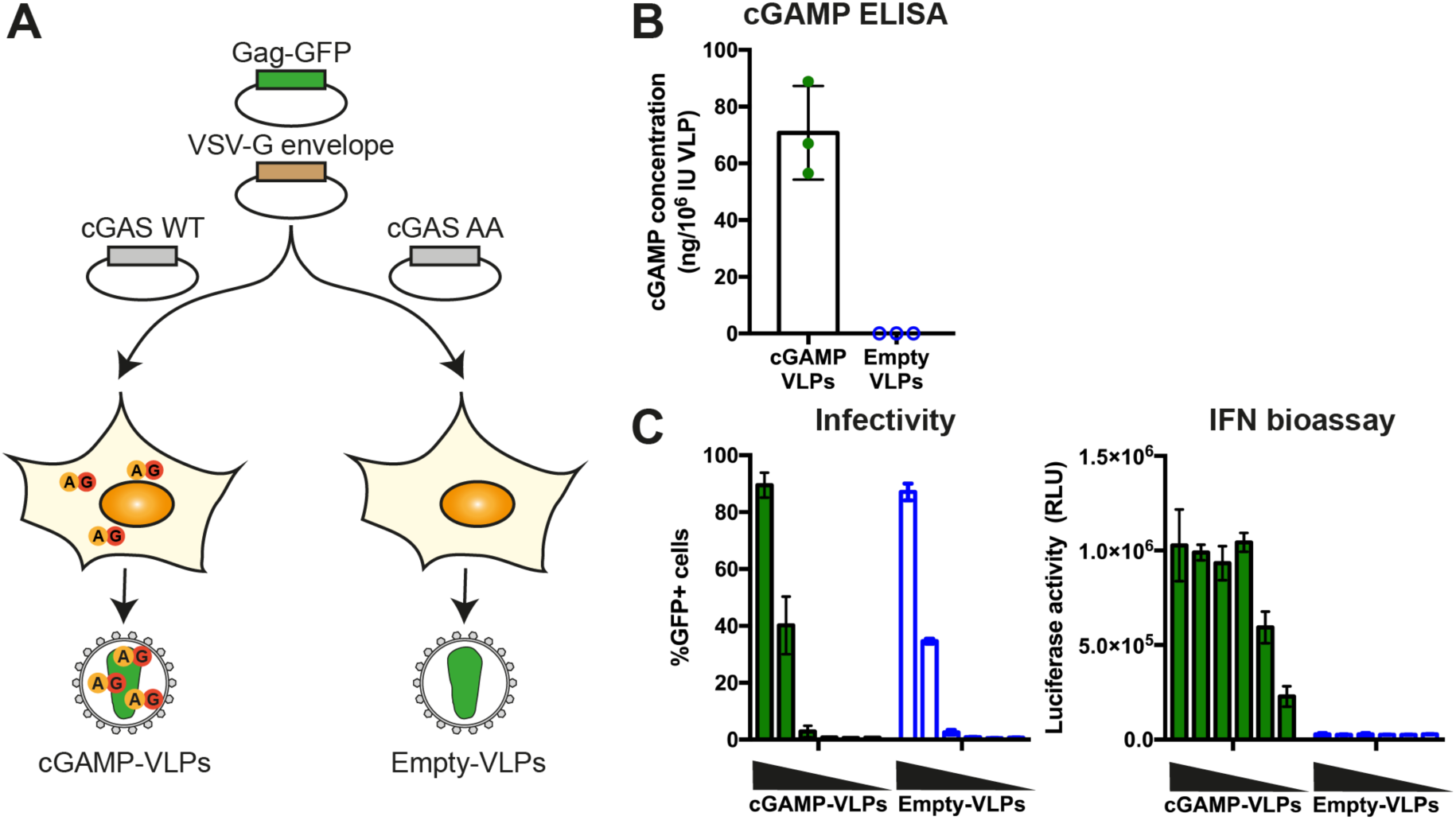
cGAMP incorporated into Gag-GFP Virus-like particles (VLPs) induces IFN-I in infected cells. **A. Schematic representation of cGAMP- and Empty-VLP production.** HEK293T cells were transfected with plasmids encoding HIV-1 Gag-GFP and VSV-G envelope to enable VLP production. Overexpression of cGAS WT in the same cells generated cGAMP that was then incorporated into nascent VLPs (cGAMP-VLPs). As control, Empty-VLPs were produced in cells where a catalytically inactive cGAS (cGAS AA) was overexpressed. **B. cGAMP is incorporated into cGAMP-VLPs.** Small molecules were extracted and the cGAMP concentration was measured using a cGAMP ELISA. **C. cGAMP-VLPs induce an IFN-I response in target cells.** HEK293 cells were infected with decreasing amounts of cGAMP-VLPs and Empty-VLPs (1/5 serial dilutions starting at 2μL of VLP stocks per well) and the infection was monitored 24 hours later by quantifying GFP^+^ cells by flow cytometry. Supernatants from the same infected cells were then transferred to a reporter cell line expressing firefly luciferase under a promoter induced by IFN-I (ISRE). Luciferase activity measured 24 hours later indicated the presence of IFN-I in the supernatants. Data in (B) are pooled from three independent VLP productions. Each symbol corresponds to one VLP production and mean and SD are shown. Data in (C) are pooled from three independent VLP productions tested simultaneously in technical duplicates in infectivity and IFN-I bioassays; mean and SD are shown.

To assess the efficiency of cGAMP incorporation into our VLPs, we extracted small molecules from VLPs as previously described (Mayer et al., 2017). cGAMP in the extract was then quantified by ELISA. While Empty-VLPs did not contain detectable levels of cGAMP, cGAMP-VLPs contained between 55 and 90 ng cGAMP per 10^6^ infectious units (IU) of VLPs (Fig 1B). We then assessed the infectivity of the VLPs and found cGAMP-VLPs and Empty-VLPs to be equally infective (Fig 1C). To confirm that cGAMP-VLPs trigger an IFN-I response, supernatant from the STING-positive HEK293 cells used in the infectivity assay was transferred to a reporter cell line expressing firefly luciferase under the interferon-sensitive response element (ISRE) promoter (Bridgeman et al., 2015). At similar infection rates, cGAMP-VLPs induced IFN-I production while Empty-VLPs did not (Fig 1C). Taken together, these results show that cGAMP can be efficiently packaged into VLPs consisting of HIV-1 Gag-GFP and the VSV-G envelope.

### Immunisation with cGAMP-VLPs induces higher and more polyfunctional CD4 and CD8 T cell responses compared to Empty-VLPs

To test whether cGAMP-VLPs induce a better immune response than Empty-VLPs *in vivo*, we injected C57BL/6 mice intramuscularly with 10^6^ infectious units of cGAMP-VLPs or Empty-VLPs or, as a control, PBS. We first assessed CD4 T cell responses in the spleen 14 days after immunisation. As we were unable to identify a specific peptide epitope within HIV Gag recognised by CD4 T cells in H-2^b^ mice, we used bone-marrow derived myeloid cells (BMMCs) pulsed with cGAMP-VLPs for evaluation of antigen-specific CD4 T cell responses. We co-cultured these cells with splenocytes for six hours before assessing IL-2, IFNγ and TNFα production by CD4 T cells by intracellular cytokine staining (ICS). Compared to mice immunised with Empty-VLPs, we observed 2.7-fold increased frequencies of CD4 T cells producing each of these cytokines in response to VLP-pulsed BMMCs in mice immunised with cGAMP-VLPs (Fig 2A, gating strategy and exemplary FACS plots in Fig S1A). Moreover, cGAMP enhanced the proportion of cells that were able to co-produce two or all three cytokines (Fig 2B; 2.1- and 3.7-fold increases, respectively).

**Fig 2:**
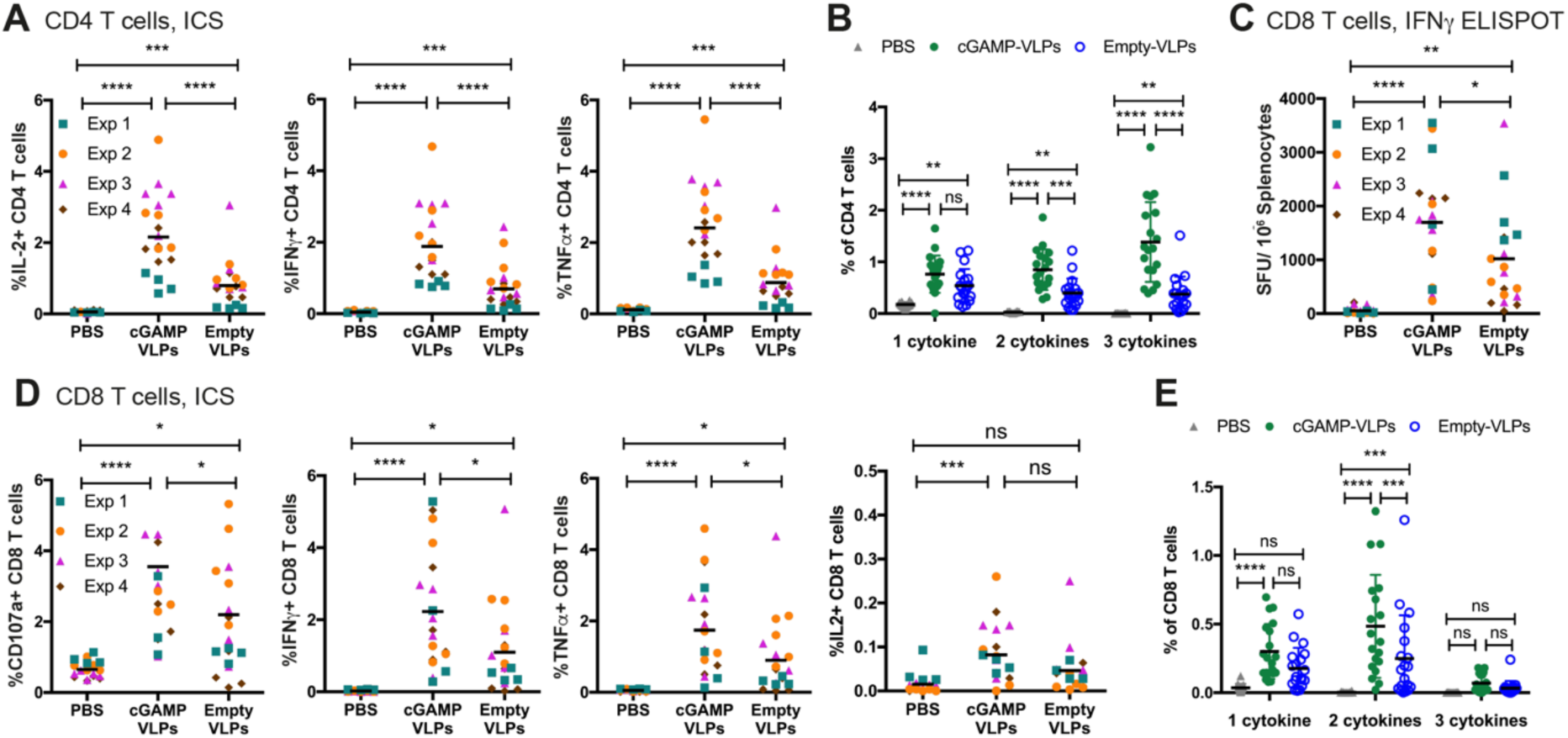
cGAMP loading of VLPs increases the magnitude of the CD4 and CD8 T cell responses elicited after immunisation. C57BL/6 mice were injected with cGAMP-VLPs, Empty-VLPs, or PBS as a control *via* the intra-muscular route. 14 days later, VLP-specific T cell responses were evaluated in the spleen. **A-B. Immunisation with cGAMP-VLPs enhances VLP-specific CD4 T cell responses.** BMMCs from C57BL/6 mice were pulsed overnight with cGAMP-VLPs and used to stimulate cells from spleens of immunised mice. Cells were co-cultured for 6 hours prior to evaluation of CD4 T cell responses by ICS. The percentage of total CD4 T cells producing each cytokine is shown in A and the percentage of CD4 T cells co-producing 1, 2 or 3 cytokines is shown in **B. C-E. Immunisation with cGAMP-VLPs facilitates induction of HIV-1 Gag-specific polyfunctional CD8 T cell responses.** Cells from spleens of immunised mice were stimulated with the HIV-SQV peptide, and responses were read out in (C) a 24-hour IFNγ ELISPOT assay; or (D, E) a 6-hour ICS assay. (D) shows the percentage of total CD8 T cells upregulating CD107a and/or producing each cytokine, and (E) shows the percentage of CD8 T cells co-producing 1, 2 or 3 cytokines. Data are pooled from four independent experiments. A total of 19 mice was analysed per condition. Symbols show data from individual animals, and in (A), (C) and (D) are colour-coded by experiment. Horizontal lines indicate the mean and SD is additionally shown in (B) and (E). Statistical analyses were performed using a 2-way ANOVA followed by Tukey’s multiple comparisons test. In (A), (C) and (D), data were blocked on experiments. ns p≥0.05; *p<0.05; **p<0.01; ***p<0.001; ****p<0.0001.

We next assessed CD8 T cell responses following immunisation. We screened a panel of overlapping 15-mer peptides spanning the HIV-1 Gag sequence and identified a peptide that stimulated an IFNγ response in cells from spleen in IFNγ ELISPOT assays (peptide p92; Fig S2A-B). We then used NetMHCpan 3.0 (http://www.cbs.dtu.dk/services/NetMHCpan-3.0/ (Nielsen and Andreatta, 2016)) to predict the optimal epitope sequence recognised within p92, and identified a 9-mer peptide (SQVTNSATI, termed HIV-SQV) that triggered T cell recognition more efficiently than the original 15-mer peptide (Fig S2C-D). This 9-mer peptide was also reported to constitute an immunodominant HIV-1 Gag epitope in H-2^b^ mice in a prior study (Holechek et al., 2016). The HIV-SQV peptide was used for all subsequent analyses of VLP-elicited CD8 T cell responses. We evaluated responses to the HIV-SQV peptide by IFNγ ELISPOT assay and showed that, when compared to Empty-VLPs, cGAMP-VLPs induced a modest but significant increase in the magnitude of the response (Fig 2C, 1.7-fold increase). To assess whether cGAMP-loading of VLPs also enhanced the polyfunctionality of the responding CD8 T cells, we stimulated splenocytes for six hours and stained for upregulation of CD107a (LAMP-1), a degranulation marker, and for the production of IFNγ, TNFα and IL-2 by ICS. Paralleling the results from the ELISPOT assay, CD8 T cells from mice immunised with cGAMP-VLPs showed a modest but significant increase in the frequency of cells upregulating CD107a (1.6-fold increase) and/or producing IFNγ (2-fold) and/or TNFα (1.9-fold) (Fig 2D, gating strategy and exemplary FACS plots in Fig S1B). Furthermore, cGAMP enhanced the proportion of CD8 T cells that were able to co-produce two of the cytokines evaluated (Fig 2E; 1.9-fold increase).

Control of vaccinia virus infection by the immune system relies in part on CD8 T cell responses (Xu et al., 2004). As immunisation with cGAMP-VLPs increased anti-HIV Gag CD8 T cell responses, we assessed whether this resulted in increased protection against subsequent infection with a vaccinia virus expressing the same HIV Gag (vVK1 (Karacostas et al., 1989)). One month after immunisation, mice were challenged with vVK1, and five days after infection virus load in the ovaries was assessed by plaque assay. We observed no weight loss over the course of the infection (Fig S3A). Immunisation with both VLPs reduced vVK1 load, and cGAMP-VLP immunised mice showed a slight but non-significant increase in protection compared to animals immunised with Empty-VLPs (Fig S3B).

Taken together, these results demonstrate that cGAMP-loading of VLPs enhances polyfunctional CD4 and CD8 T cell responses to VLP antigens.

### cGAMP loading of VLPs enhances serum titres of VLP binding and neutralising antibodies

Next, we assessed the antibody response in immunised mice. We set up ELISAs that allow detection of serum antibodies binding to any protein in the VLPs, or of antibodies specific for the VSV-G envelope or the HIV-Gag protein. In mice immunised with VLPs, we detected very strong IgG responses and lower-titre IgM responses targeting the total VLP protein pool 14 days after immunisation, indicating antibody class-switching (Fig 3A and Fig S4A). Interestingly, immunisation with cGAMP-VLPs induced stronger anti-VLP antibody responses compared to the Empty-VLP immunised group, with statistically significant differences being observed in IgG2a/c, IgG2b and IgM levels. We also detected IgG antibodies targeting the VSV-G envelope, and IgG1, IgG2a/c and IgG2b titres were higher in the cGAMP-VLP immunised group (Fig 3B). Titres of antibodies recognising the intracellular antigen HIV-Gag were low or undetectable, but a similar trend was observed for a higher-magnitude response in the cGAMP-VLP immunised group (Fig 3C and S4A).

**Fig 3:**
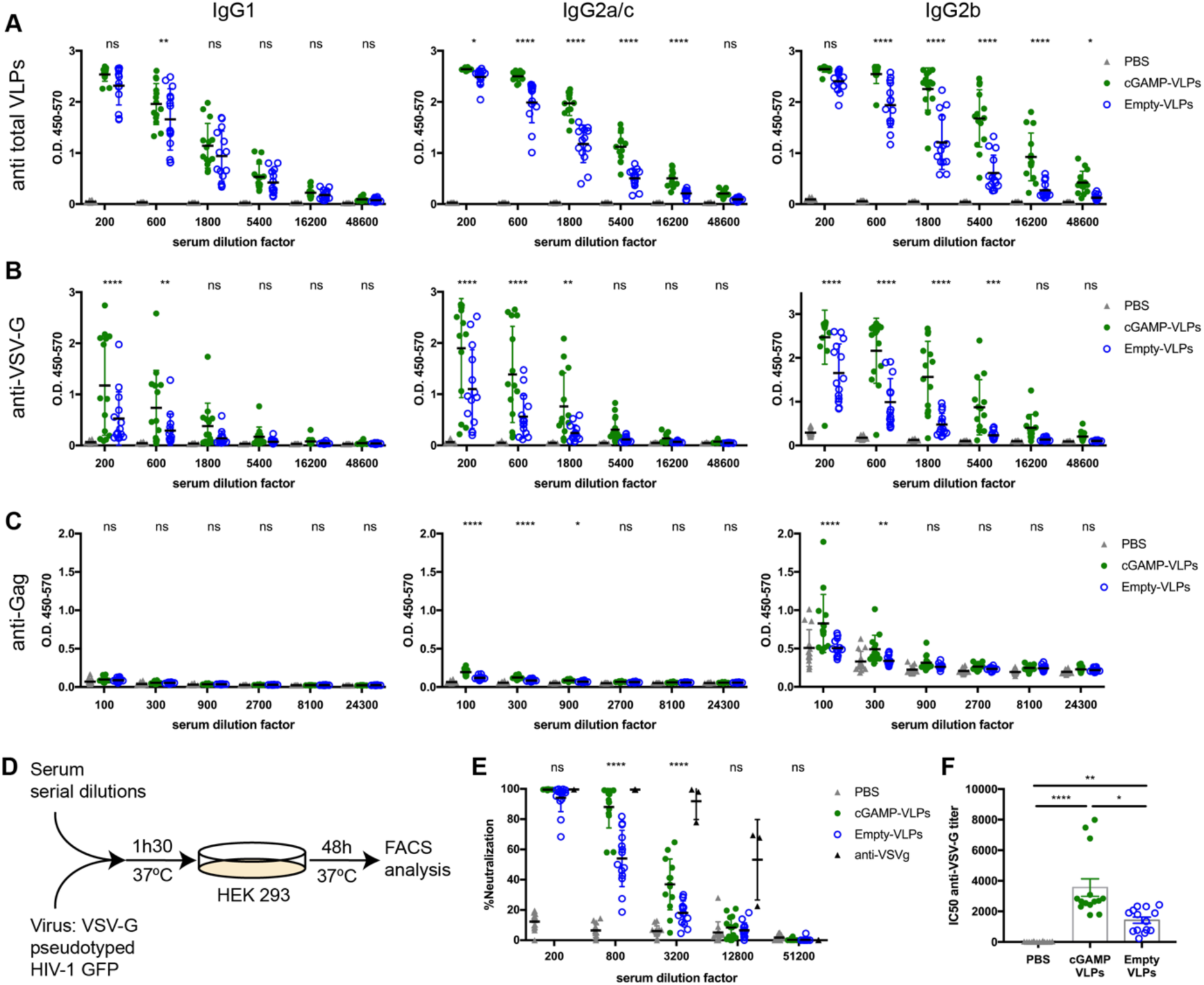
Immunisation with VLPs containing cGAMP increases neutralising antibody responses. C57BL/6 mice were injected with cGAMP-VLPs, Empty-VLPs, or PBS as a control *via* the intramuscular route. Serum antibody responses were evaluated 14 days later. **A-C. cGAMP loading enhances IgG responses specific to VLP proteins, including VSV**-**G.** ELISA plates were coated with lysate from cGAMP-VLPs (A), recombinant VSV-G protein (B), or recombinant HIV-1 Gag protein (C). Antibodies of different isotypes specific for these proteins were measured in sera from immunised mice by ELISA. The optical density at increasing serum dilutions is shown. **D-F. Immunisation with cGAMP-VLPs enhances production of neutralising antibodies.** Serial dilutions of serum samples from individual mice were incubated with VSV-G pseudotyped HIV-1-GFP for 90 minutes at 37°C before infection of HEK293 cells. As a control, serial dilutions of the anti-VSV-G neutralising antibody 8G5F11 were tested in parallel. After two days, infection was measured by quantifying GFP^+^ cells by flow cytometry (D). Neutralising capacities of serum samples from individual animals were calculated as a percentage of neutralisation (calculated relative to the maximum infection in each experiment) (E) and as the half maximal inhibitory concentration (IC50) (F). Data are pooled from three independent experiments. A total of 14 mice was analysed per condition. Symbols show data from individual animals, and the mean and SD are indicated. Statistical analyses were done using a 2-way ANOVA followed by Tukey’s multiple comparisons test, only showing significance between cGAMP-VLPs and Empty-VLPs (A-E) or a Kruskall-Wallis test followed by Dunn’s multiple comparisons test (F). ns p≥0.05; *p<0.05; **p<0.01; ***p<0.001; ****p<0.0001.

To test whether the anti-VLP antibodies were neutralising, we assessed the *in vitro* neutralisation capacity of sera using a VSV-G pseudotyped HIV-1-based lentivector expressing GFP (Fig 3D). The effect of pre-incubation with serum samples on the infectivity of the HIV-1-GFP virus was measured by monitoring GFP expression in HEK293 cells (Fig S4B-C). Although immunisation with both cGAMP-VLPs and Empty-VLPs induced neutralising antibodies, this response was stronger when cGAMP was present within the VLPs, and sera from cGAMP-VLP immunised mice showed a 2.5-times higher half maximal inhibitory concentration (Fig 3E-F). In summary, immunisation with cGAMP-VLPs induced an increased antibody response that targeted proteins from total VLP lysates including the VSV-G envelope protein. Moreover, cGAMP-loading enhanced production of virus neutralising antibodies.

### Incorporation of cGAMP into VLPs increases the CD4 Tfh cell response

To gain insight into how immunisation with cGAMP-VLPs resulted in an increased antibody response, we investigated B and T cell populations in inguinal lymph nodes that drain the injection site. As CD4 T cell responses were increased in the spleens of cGAMP-VLP immunised mice, we first tested whether follicular CD4 T cell numbers were elevated in lymphoid tissues draining the immunisation site. We identified follicular CD4 T cells as CD4^+^CD44^+^PD1^hi^CXCR5^hi^ and subdivided them into Tfh and Tfr cells by analysing FoxP3, which is expressed in Tfr cells (Fig 4A). Immunisation with VLPs led to an increase in the proportion of follicular T cells within the CD4 T cell population in the draining lymph node (Fig 4A-B). This was due to an expansion of Tfh cells, as the latter increased significantly in frequency after VLP immunisation, whereas Tfr frequencies within CD4 T cells remained unaltered. As a consequence of this, there was a profound shift in the Tfh:Tfr ratio in VLP-immunised as compared to control mice (Fig 4C). Importantly, the increase in Tfh cells was more pronounced in cGAMP-VLP immunised mice compared to Empty-VLP injected animals (Fig 4B-C; 1.6-fold increase).

**Fig 4:**
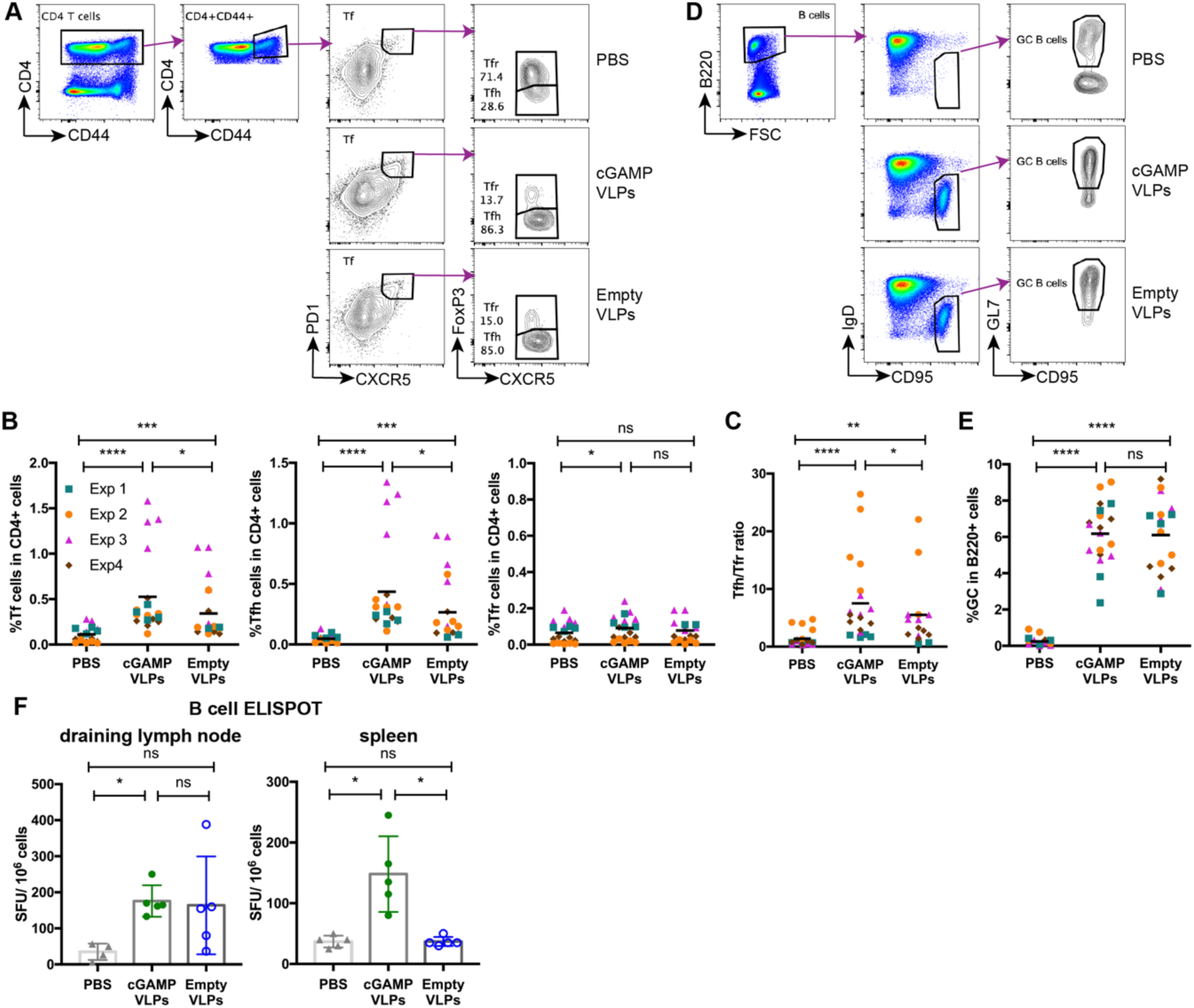
cGAMP loading of VLPs enhances induction of CD4 Tfh responses. C57BL/6 mice were injected with cGAMP-VLPs, Empty-VLPs, or PBS as a control *via* the intra-muscular route. 14 days later, T and B cells in the draining inguinal lymph nodes were characterised by flow cytometry and B cell ELISPOT assays. **A-C. Immunisation with cGAMP-VLPs enhances accumulation of Tfh cells in the draining lymph node.** T follicular (Tf) cells were identified by flow cytometry as CD4^+^CD44^+^CXCR5^hi^PD1^hi^ cells and were further subdivided into Tfr cells (FoxP3^+^) and Tfh cells (FoxP3^-^). The gating strategy is shown in (A) and the percentages of Tf, Tfh and Tfr cells within CD4^+^ cells are shown in (B). The ratio of Tfh/Tfr is shown in C. **D-E. Immunisation with VLPs induces germinal centre formation.** Germinal centre B cells were identified by flow cytometry as B220^+^IgD^-^CD95^+^GL7^+^ cells. The gating strategy is shown in (D) and the percentage of germinal centre B cells amongst B220^+^ cells is shown in (E). **F. Immunisatio nwith cGAMP-VLPs increases production of antibody-secreting cells.** Cells from draining lymph nodes and spleens were seeded in ELISPOT plates coated with VLP lysates. After overnight incubation, cells producing VLP-specific IgG antibodies were identified using an anti-IgG Fc antibody. In (B), (C) and (E), data were pooled from four independent experiments including a total of 19 mice analysed per condition. Symbols show data from individual animals and are colour-coded by experiment. Horizontal lines indicate the mean. In (F), symbols show data from 5 mice per group measured in duplicate in one experiment. Mean and SD are indicated. Statistical analyses were done using a 2-way ANOVA followed by Tukey’s multiple comparisons test (B, C, E) or a Kruskall-Wallis test followed by Dunn’s multiple comparisons test (F). ns p≥0.05; *p<0.05; **p<0.01; ***p<0.001; ****p<0.0001.

To assess the impact of this increased Tfh response on B cell responses, we first gated on GC B cells (B220^+^IgD^-^CD95^+^GL7^+^ cells; Fig 4D). Immunisation with VLPs induced a robust GC B cell response, with no difference being observed in the frequencies of GC B cells in cGAMP-VLP and Empty-VLP groups at the day 14 time-point analysed (Fig 4E). We next evaluated the generation of antibody-secreting cells (ASCs) by antigen-specific B cell ELISPOT assay on cells from both draining lymph nodes and spleens 14 days after immunisation. VLP-specific ASCs were detected in the lymph nodes of mice injected with both cGAMP-VLPs and Empty-VLPs (Fig 4F). In the spleen, VLP-specific ASCs were also observed in cGAMP-VLP immunised animals, but not in Empty-VLP immunised animals (Fig 4F). Taken together, these results suggest that immunisation with cGAMP-VLPs increased the antibody response by enhancing the accumulation of Tfh cells in draining lymph nodes, thereby promoting the development of ASCs.

### cGAMP-VLPs pseudotyped with IAV haemagglutinin induce a neutralising antibody response and confer protection following live virus challenge

As immunisation with cGAMP-VLPs induced high titres of neutralising antibodies, we explored whether they could confer protection following a live virus challenge. Protection against IAV infection correlates with serum antibody responses against the surface glycoprotein, haemagglutinin (HA) (Krammer, 2019). We therefore produced cGAMP-VLPs and Empty-VLPs incorporating HA from the mouse-adapted PR8 strain of IAV (designated cGAMP-HA-VLPs and Empty-HA-VLPs, respectively) (Fig S5A). cGAMP-HA-VLPs and Empty-HA-VLPs were equally infective, as assessed by the percentage of GFP-expressing cells observed following infection of HEK293 cells with titrated doses of VLPs (Fig S5B). Staining of infected HEK293 cells with an antibody recognising HA revealed the presence of similar percentages of HA^+^ cells after infection with cGAMP-HA-VLPs and Empty-HA-VLPs (Fig S5B). As control, cells infected with cGAMP-VLPs without HA showed no detectable staining. These data confirmed that HA was transferred by HA-VLPs to infected cells. Finally, we verified that supernatant from cells infected with cGAMP-HA-VLPs contained IFN-I, suggesting that the presence of HA did not affect the incorporation of cGAMP into the VLPs (Fig S5C).

Next, we immunised mice with HA-VLPs. Two weeks after immunisation, sera were analysed for neutralising antibodies using a micro-neutralisation assay. In brief, a single cycle IAV expressing eGFP and PR8 HA was pre-incubated with sera and its infectivity was then monitored using MDCK-SIAT1 cells (Powell et al., 2012). Immunisation with both Empty-HA-VLPs and cGAMP-HA-VLPs induced neutralising antibodies, and the presence of cGAMP in the VLPs increased this response by 2.7-fold (Fig 5A). To determine if immunisation conferred protection upon *in vivo* challenge with live IAV, we infected mice with 10^4^ TCID50 of HA-matched PR8 IAV one month after immunisation. Animals immunised with 10^6^ infectious units of both VLPs were protected against the weight loss observed between day three and four after IAV infection in PBS-treated mice, both resulting in 100% survival (Fig 5B-C). This prompted us to reduce the amount of VLPs used for immunisation. At an intermediate dose of 2×10^5^ infectious units of VLPs, cGAMP-HA-VLPs were fully protective against weight loss and disease progression to an endpoint where humane sacrifice was necessary, while immunisation with Empty-HA-VLPs delayed disease progression by about four days, resulting in 100% and 16.7% survival, respectively (Fig 5B-C). At the lowest dose of VLPs tested (5×10^4^ infectious units), Empty-HA-VLPs were not protective whereas cGAMP-HA-VLPs protected most animals against severe disease (83% survival) (Fig 5B-C).

**Fig 5:**
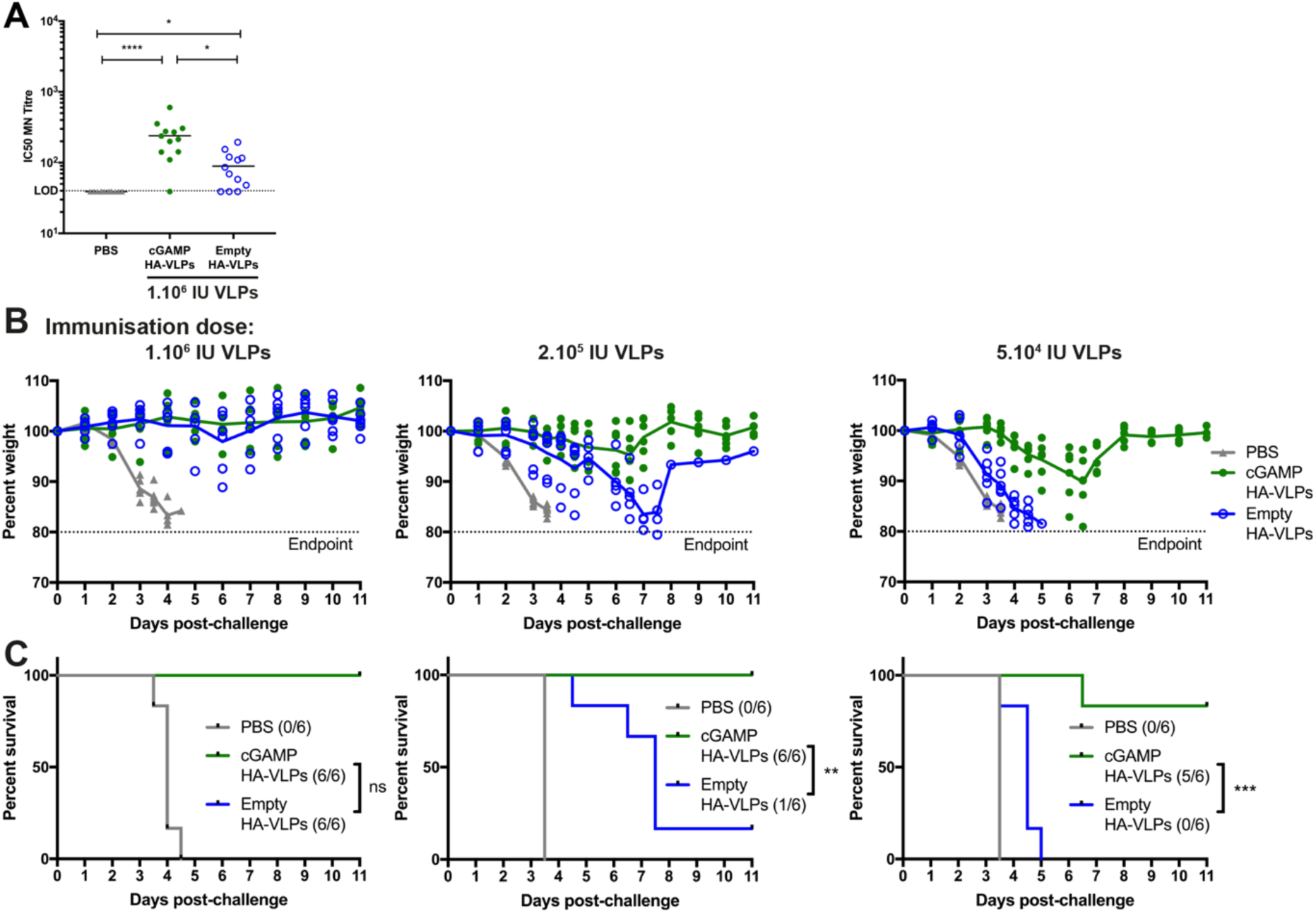
cGAMP-VLPs pseudotyped with IAV HA induce neutralising antibodies and confer protection following IAV infection. Mice were immunised with PBS as a control, cGAMP-HA-VLPs or Empty-HA-VLPs *via* the intra-muscular route. **A. VLPs pseudotyped with IAV HA induce neutralising antibodies.** Two weeks after immunisation with 10^6^ infectious units of VLPs, sera were collected, heat-inactivated, and titres of antibodies capable of neutralising an IAV expressing a matched HA protein were determined by micro-neutralisation (MN) assay. The dotted line shows the limit of detection (LOD). **B-C. Low doses of cGAMP-HA-VLPs are able to confer protection following IAV challenge.** One month after immunisation with the indicated doses of VLPs, animals were infected with 10^4^ TCID_50_ of IAV PR8 virus. Weight loss was monitored over the following eleven days and is shown as a percentage of starting weight (B). Animals approaching the humane end-point of 20% weight loss were culled and survival to end-point curves are shown in (C). In (A), data were pooled from two independent experiments including a total of 10 mice per condition. In (B) and (C), 5 mice per group were analysed for each VLP dose. In (A) and (B), symbols show data from individual animals. Statistical analyses were done using a Kruskall-Wallis test followed by Dunn’s multiple comparisons test (A) or a survival analysis with the Log-rank (Mantel-Cox) test (C). ns p≥0.05; *p<0.05; **p<0.01; ***p<0.001; ****p<0.0001.

Taken together, these results show that vaccination with VLPs incorporating IAV HA induced neutralising antibodies in mice, which were protected against subsequent IAV challenge. The presence of cGAMP in HA-VLPs enhanced the antibody response and, at lower doses of VLPs used for immunisation, facilitated protection against IAV.

## Discussion

New and more targeted adjuvants are needed for improved efficacy and safety of vaccines. There is growing interest in using adjuvants that specifically activate innate immune pathways used by cells to detect viral infections. cGAMP is one such example. cGAMP is a natural molecule produced by cells upon virus infection that specifically triggers STING and thereby induces innate and adaptive immune responses. The vaccination strategy we describe here is based on coupling the adjuvant cGAMP with antigen(s) in a single entity, namely HIV-derived VLPs. We demonstrate that cGAMP-loading of these VLPs increased CD4 and CD8 T cell responses, as well as antibody responses, against protein antigens in the VLPs. Furthermore, vaccination with VLPs containing cGAMP protected mice against disease development following infection with a virus expressing a cognate antigen.

HIV-derived VLPs are a flexible system that allows incorporation of proteins of choice. We demonstrate this by decorating VLPs with IAV HA, and show that upon immunisation, these VLPs induced antibodies that neutralised IAV expressing a matched HA protein. In future, other pathogen-derived proteins could be incorporated into cGAMP-loaded VLPs as a strategy to produce vaccines for a diverse breadth of pathogens. For example, multiple HA proteins from different IAV clades could be incorporated to induce broadly protective responses or envelope proteins from other viruses such as Zika or Ebola could be delivered using this approach.

The VLPs used here were pseudotyped with VSV-G, which has a broad tropism. It is possible to replace VSV-G with other envelope proteins that target VLPs to specific cell types. For example, the envelope protein from Sindbis virus or antibodies such as those to DEC205 target virus particles to DCs, an essential antigen presenting cell type (Trumpfheller et al., 2006; Yang et al., 2008). It will be interesting to determine whether DC-targeted VLPs containing cGAMP have a similar effect on the responses induced compared to the VSV-G pseudotyped VLPs described here. DC targeting could improve vaccine safety by restricting cGAMP delivery to relevant antigen-presenting cells, thereby limiting systemic inflammation. Indeed, with the VSV-G pseudotyped VLPs used here, we observed a transient weight loss of approximately 5% in cGAMP-VLP but not in Empty-VLP immunised mice (data not shown).

Incorporating cGAMP inside viral particles likely increases its stability at the site of injection by preventing degradation in the extracellular milieu. In addition to HIV-derived lentiviruses, other enveloped viruses also incorporate cGAMP (Bridgeman et al., 2015; Gentili et al., 2015). Therefore, our strategy of protecting the adjuvant cGAMP together with antigen in viral particles may be applicable to other viral vectored vaccines such as modified vaccina virus Ankara (MVA).

Many studies are currently aimed at designing vaccines that induce antigen-specific CD8 T cells (Panagioti et al., 2018). We found that cGAMP-loading of VLPs modestly enhanced CD8 T cell responses to the internal HIV-Gag antigen. The increased response in cGAMP-VLP immunised mice did not result in a significant improvement in protection against a vaccinia virus expressing the same HIV-Gag compared to that observed in animals vaccinated with Empty-VLPs. However, the immunisation strategy employed here consisted of a single dose of VLPs and may be improved by employing prime-boost strategies. In light of the neutralising antibody response induced by cGAMP-loaded VLPs, heterologous booster immunisations using non-particulate vaccines or viral particles with a different envelope protein are particularly promising.

Both splenic CD4 effector T cell responses as well as Tfh cell numbers in draining lymphoid tissues were enhanced by incorporation of cGAMP in VLPs. It is likely that these effects explain the increased antibody responses we observed against VLP proteins. The CD4 T cell response was skewed toward a Th1 phenotype, as indicated by robust IFNγ and TNFα production by CD4 T cells and enhanced IgG2a/c and IgG2b antibody responses. Both the cell type mediating antigen presentation as well as the cytokines produced at the time of T cell activation are crucial for polarisation of T cell responses (Hong et al., 2018; Itano and Jenkins, 2003; O’Garra, 1998). Notably, IFN-I and IL-6 production by DCs have been reported to induce the development of Tfh cells in mice (Cucak et al., 2009; Nurieva et al., 2009; Riteau et al., 2016).We previously found that cGAMP-loaded viruses induce IFN-I in bone-marrow derived macrophages *in vitro* (Bridgeman et al., 2015). Activation of STING and down-stream IRF3 and NF-κB signalling by cGAMP *in vivo* might therefore trigger production of IFN-I and IL-6 that could underlie the potent CD4 Tfh response elicited following immunisation with cGAMP-VLPs. The specific cell types infected by VLPs *in vivo* and the cytokines induced by these cells are likely to be key aspects of the response induced by cGAMP-VLPs *in vivo* and warrant further investigation.

VLPs bearing IAV HA induced a neutralising antibody response and protected immunised mice against development of severe disease following challenge with live IAV. Importantly, the presence of cGAMP in VLPs enabled induction of a protective response even at low VLP doses. cGAMP-loading of viral vectored vaccines may therefore allow the vaccine dose administered to be reduced without compromising vaccine efficacy. We believe this will be advantageous in at least two ways: by increasing safety and by reducing cost of vaccine production. The latter is particularly important for lentivirus-based vectors that can typically only be produced at lower titres than other viral vectored vaccines.

In summary, we provide evidence that vaccination with HIV-derived VLPs containing both the adjuvant cGAMP and protein antigens constitutes an efficacious platform for induction of CD8 T cell and neutralising antibody responses. This VLP-based strategy of coupling adjuvant and antigen in a single entity is therefore a promising approach for development of new and safer vaccines against a range of pathogens.

## Materials and methods

### Mice

All mice were on the C57Bl/6 background. This work was performed in accordance with the UK Animals (Scientific Procedures) Act 1986 and institutional guidelines for animal care. This work was approved by project licenses granted by the UK Home Office (PPL No. 40/3583, No. PC041D0AB and No. PBA43A2E4) and was also approved by the Institutional Animal Ethics Committee Review Board at the University of Oxford.

### Cells

Cell lines (HEK293T, HEK293, 3C11, 143B, MDCK-SIAT1, MDCK-PR8) were maintained in DMEM (Sigma Aldrich) supplemented with 10% FCS (Sigma Aldrich) and 2mM L-Glutamine (Gibco) at 37°C and 5% CO_2_. 3C11 cells are HEK293 cells stably transduced with an ISRE-Luc reporter construct (Bridgeman et al., 2015). 143B cells were a kind gift from N. Proudfoot (University of Oxford).

Bone marrow cells were isolated from humanely killed adult mice by standard protocols and grown in 6-well plates for 5 days in RPMI supplemented with 10% FCS, 2mM L-Glutamine, 1% PenStrep and 20ng/mL mouse GM-CSF to obtain bone marrow-derived myeloid cells (BMMCs).

### Reagents and antibodies

See Supplementary Table 1

### VLP and HIV-1 vector production

All VLPs were produced by transient transfection of HEK293T cells with Fugene 6 (Promega, ref E2691). HEK293T were seeded in 15-cm dishes to reach 60-70% confluency the next day and VLPs were produced by co-transfecting plasmids encoding Gag-eGFP and the VSV-G envelope (pGag-EGFP and pCMV-VSV-G, respectively) at a ratio of 2:1. VLPs were loaded with cGAMP by co-transfecting at the same time a plasmid encoding mouse cGAS WT (pcDNA3-Flag-mcGAS). Empty-VLPs were produced as control by co-transfecting a catalytically inactive mouse cGAS (cGAS AA; pcDNA3-Flag-mcGAS-G198A/S199A). One day after transfection, the medium was changed. Supernatants were collected 24, 32 and 48 hours after medium change, centrifuged and filtered (Cellulose Acetate membrane 0.45 μm pore-size). At each media change VLPs were concentrated by ultracentrifugation through a 20% sucrose cushion at 90,000g for 2.5 hours at 8°C using a Beckman SW32 rotor. VLPs were resuspended in PBS and subsequent harvests were resuspended using the resuspended VLPs from previous harvests to maximise titre.

For pseudotyping cGAMP-VLPs and Empty-VLPs with Influenza Haemagglutinin H1 (HA; pcDNA3.1-H1 (PR8)), cells were transfected as above with the following plasmids: Gag-eGFP, VSV-G, HA and cGAS WT or AA at a ratio of 2:1:1:2.

To produce VSV-G pseudotyped HIV-1 vectors for neutralisation assays, HEK293T cells were co-transfected with the following plasmids: HIV-1 NL4-3 *Δ*Env GFP (pNL4-3-deltaE-EGFP) and VSV-G at a ratio of 2:1.

### VLP Titration and cGAMP incorporation assays

HEK293 cells were seeded at a density of 1×10^5^ cells per well in 24-well plates. The next day, cells were infected with decreasing amounts of VLPs in the presence of 8μg/mL of polybrene. 24 hours after infection, cells were collected and first stained with anti-CD16/32 and Aqua fixable Live/Dead in FACS Buffer (PBS, 1% FCS, 2mM EDTA) for 15 minutes at RT. Cells used for titration of HA-VLPs were also stained for HA using a primary human anti-H1 & H5 antibody in FACS Buffer for 30 minutes at 4°C. Cells were then washed twice and further stained with a secondary goat anti-human Alexa Fluor 647-conjugated antibody for 30 minutes at 4°C, followed by two washes. All cells were fixed using BD Cellfix before acquisition on an Attune Nxt flow cytometer. Infection was measured by analysing GFP positive cells by flow cytometry using FlowJo version 10. VLP titres were calculated based on the number of GFP^+^ cells compared to the number of cells in the well at the time of infection and expressed as infectious units/mL (IU/mL). Supernatants from infected cells were transferred onto ISRE reporter cells to assess IFN-I production in response to cGAMP incorporated in VLPs as described previously (Bridgeman et al., 2015). After 24 hours of incubation with supernatants, expression of the ISRE-Luc reporter was assessed using the One-Glo luciferase assay system. Small molecular extracts were prepared from VLPs as described (Mayer et al., 2017). Briefly, 2×10^6^ IU of ultra-centrifuged VLPs resuspended in PBS were lysed in X-100 Buffer (1mM NaCl, 3mM MgCl2, 1mM EDTA, 1% Triton X-100, 10mM Tris pH7.4) by adding 1/10 volume of 10X buffer for 20 minutes on ice while vortexing regularly. After centrifugation at 1,000g for 10 minutes at 4°C, supernatants were treated with 50U/mL of benzonase for 45 minutes on ice. Samples were then extracted with phenol-chloroform, and the aqueous phase was then transferred to Amicon Ultra 3K filter columns. After filtering by centrifugation at 14,000g for 30 minutes at 4°C, samples were dried in a SpeedVac and resuspended in 200μL of water. cGAMP was quantified using the 2’-3’ cGAMP ELISA kit following manufacturer’s instructions.

### IAV

The Influenza virus H1N1 A/Puerto Rico/8/1934 (Cambridge) (PR8) and the non-replicating S-FLU vector expressing eGFP (S-eGFP) were generated as previously described (Powell et al., 2012). Plasmids encoding the IAV Cambridge strain of A/Puerto Rico/8/34 were used to generate the wild type H1N1 A/Puerto Rico/8/1934 (Cambridge) (PR8) seed virus. The same plasmids were used to generate the PR8 S-eGFP with slight modifications: the HA coding region in the plasmid expressing HA viral RNA was replaced with eGFP and an additional plasmid was included to provide a functional PR8 HA in *trans* to rescue the PR8 S-eGFP seed virus. Briefly, the plasmids were transfected into HEK293T cells using lipofectamine 2000 and supernatant containing seed virus was collected 72 hours after transfection. The wild type PR8 virus and the S-eGFP vector were then propagated by infecting MDCK-SIAT1 cells or MDCK-SIAT1 stably transfected with PR8 HA (MDCK-PR8), respectively, with seed virus, followed by medium change into VGM (Viral Growth Media; DMEM, 1% BSA, 10mM HEPES buffer, 1% PenStrep) containing 1µg/mL TPCK-treated trypsin. Viruses were harvested 48 hours later. The TCID50 was determined by infecting MDCK-SIAT1 or MDCK-PR8 cells with a ½-log dilution series of viruses in VGM for one hour in eight replicates using 96-well flat-bottom plates. Next, 150µl per well of VGM with TPCK-treated trypsin (1 µg/mL) was added and cells were further incubated for 48 hours at 37°C. The PR8 virus and the S-eGFP vector were quantified by Nucleoprotein (NP) staining and eGFP expression respectively and TCID50 was calculated using the method of Reed and Muench (Reed and Muench, 1938).

### Vaccinia virus

Stocks of the vaccinia virus expressing HIV-1 HXB.2 Gag (vVK1) were produced by growth in 143TK-cells, and infectious virus titers were determined by plaque assay (Borrow et al., 1994).

### Immunisation and viral challenge of mice

C57Bl/6 female mice between 6-8 weeks old were obtained from University of Oxford Biomedical Services or Envigo RMS (UK) Limited. Animals were injected intra-muscularly with 50μL per hindleg of PBS or 10^6^ IU of cGAMP-VLPs or Empty-VLPs, unless otherwise stated, under inhalation isoflurane (IsoFlo, Abbott) anaesthesia. Weight was monitored every day for 14 days. For immunophenotyping, mice were culled on day 14 by inhalation of carbon dioxide and cervical dislocation. For viral challenge experiments, mice were monitored every other day for an additional two weeks before challenge.

For IAV challenge, blood samples were acquired two weeks after immunisation for evaluation of the serum antibody response. Mice were then challenged a month after immunisation *via* the intranasal route with 10,000 TCID_50_ of PR8 diluted in 50μL VGM under inhalation isoflurane anaesthesia. Weight was monitored daily and mice were culled by inhalation of carbon dioxide and cervical dislocation when body weight loss approached the humane end-point of 20%.

For vaccinia virus challenge, mice were infected *via* the intra-peritoneal route with 10^6^ PFU vVK1 in 100μL PBS. Weight was monitored daily for five days. Animals were then culled by inhalation of carbon dioxide and cervical dislocation, and ovaries were collected for virus titration.

### Analysis of T cell responses by ICS and ELISPOT

Splenocytes were obtained by separating spleens through a 70μm strainer, and were then treated with red blood cell lysis buffer for 5 minutes, washed and resuspended in RPMI supplemented with 2% Human serum, 2mM L-Glutamine, 1% PenStrep (R2).

For ELISPOT assays, splenocytes were seeded in R2 at a density of 1.5×10^5^ cells per well on ELISPOT plates pre-coated with anti-IFNγ detection antibody. Cells were either non-treated or treated with 2μg/mL HIV-1 Gag peptide or with 10ng/mL PMA and 1μg/mL ionomycin as a control, and incubated for 48 hours at 37°C before detection according to the manufacturer’s instructions (Mouse IFNγ ELISPOT BASIC (ALP) kit).

For intracellular cytokine staining (ICS), cells were seeded in R2 at a density of 1×10^6^ cells per well in a round-bottom 96 well plates. Cells were either non-treated or treated with 2μg/mL HIV-SQV 9-mer peptide or co-cultured with BMMCs pulsed overnight with cGAMP-VLP at a multiplicity of infection of 1. Cells were also treated with 10ng/mL PMA and 1μg/mL ionomycin as a positive control. After 1 hour of incubation at 37°C, Golgi STOP was added according to manufacturer’s instructions. After a further 5 hours of incubation at 37°C, cells were washed twice in FACS buffer (PBS, 1% FCS, 2mM EDTA), incubated with anti-CD16/32 and Aqua or violet fixable Live/Dead in FACS Buffer for 15 minutes at RT and were then washed twice in FACS Buffer. Subsequent extracellular staining involved incubation of cells for 30 minutes at 4°C with the following antibodies: anti-CD8 BV605 and anti-CD90.2 PerCP-Cy5.5 in FACS Buffer for CD8 T cells analysis in cells stimulated with the HIV peptide, or anti-CD4 AF700, anti-CD8 BV605 and anti-MHC-II BV510 in Brilliant stain buffer for CD4 T cells analysis in cells stimulated with pulsed BMMCs. Cells were then washed twice in FACS Buffer and fixed using BD Cytofix/Cytoperm buffer for 20 minutes at 4°C. After 2 washes in FACS Buffer with 10% BD Cytoperm/wash, intracellular staining was performed for using anti-TNFα PE, anti-IFNγ PE-Cy7 and anti-IL2 APC in FACS Buffer with 10% BD Cytoperm/wash for 30 minutes at 4°C. After 2 washes in FACS Buffer with 10% BD Cytoperm/wash, cells were fixed for 10 minutes at RT in BD Cellfix, washed again and resuspended in FACS Buffer for acquisition on Attune NxT flow cytometers. Analysis was performed using FlowJo version 10. Gates for phenotypic markers of CD4 and CD8 T cells were based on FMO controls. Unstimulated control cells were used for other gates.

### Analysis of serum antibody titres by ELISA

To extract protein, VLPs were lysed in PBS containing 0.5% Triton X-100 and 0.02% Sodium Azide for 10 minutes at RT. Quantity of protein extracted was quantified by BCA assay. Costar high-binding half-area flat bottom 96 well plates were coated overnight at 4°C with either 10μg/mL cGAMP-VLP lysates, 0.5μg/mL recombinant HIV-1 IIIB pr55 Gag protein or 1.5μg/mL recombinant VSV-G protein. The next day, plates were washed twice in PBS, then twice in PBS with 0.1% Tween-20 (wash buffer) and blocked in PBS with 3% BSA for 2 hours at RT. Sera collected on day 14 after immunisation were serially diluted in PBS with 0.5% BSA starting at a dilution of 1/200 and diluting 1/3. After four washes in wash buffer, serum dilutions were added to the plates in duplicates (25μl per well) and incubated for 1 hour at 37°C. Plates were washed four times in wash buffer. Next, 50μl per well of HRP-conjugated antibodies recognising different antibody classes or subclasses were added using the following dilutions: goat anti-mouse IgG1 / IgG2a/c / IgG2b, 1/10,000; IgM, 1/2,000 in PBS with 0.5% BSA and incubated for 1 hour at RT. Plates were washed four times in wash buffer and 50μl of TMB substrate was added per well. Plates were incubated for approximately 30 minutes or until the signal was saturating and 50μl of STOP solution was added per well before reading absorbance at 450nm and 570nm.

### Analysis of germinal centre B cells and T follicular cells in draining lymph nodes

Both inguinal lymph nodes were meshed through a 70μm strainer. 10^6^ cells per animal were used for each staining.

For germinal centre B cell analysis, cells were first stained with anti-CD16/32 and Aqua fixable Live/Dead in FACS Buffer for 15 minutes at RT. After two washes in FACS Buffer, extracellular staining was performed using anti-B220 APC-Cy7, anti-CD95 PE, anti-IgD PerCP-Cy5.5 and GL7 AF647 in FACS Buffer for 30 minutes at 4°C. Cells were then washed, fixed for 10 minutes at RT in BD Cellfix, washed again and resuspended in FACS Buffer.

For T follicular cell analysis, cells were first stained with anti-CD16/32 and Aqua fixable Live/Dead in FACS Buffer for 15 minutes at RT. After two washes in FACS Buffer, extracellular staining was performed using anti-B220 BV510, anti-CD4 AF700, anti-CD44 PerCP-Cy5.5, anti-CXCR5 BV421 and anti-PD-1 APC in Brilliant stain buffer for 1 hour at 4°C. After two washes in FACS Buffer, cells were fixed using the eBioscience FoxP3 fixation buffer for 25 minutes at RT. Cells were then washed twice in cold eBioscience Perm buffer and intracellular staining was performed using anti-FoxP3 PE-Cy7 in eBioscience Perm buffer for 40 minutes at RT. After two washes in eBioscience Perm buffer, cells were then fixed for 10 minutes at RT in BD Cellfix, washed again and resuspended in FACS Buffer for acquisition on Attune NxT flow cytometers. Analysis was performed using FlowJo version 10. Gates for phenotypic markers of CD4 T cells and B cells were based on FMO controls, and gates for GC and Tfh/Tfr markers were based on PBS-immunised mice.

### B cell ELISPOT

Cells from spleen and draining lymph nodes were collected as described above and counted. Three different amounts of cells (10^6^, 3×10^5^, 1×10^5^) were seeded in duplicate in R2 on ELISPOT plates coated overnight with 2μg/mL of lysates from cGAMP-VLPs (see ELISA). Plates were then incubated overnight at 37°C before detection according to manufacturer’s instruction (Mouse IgG Basic ELISPOT BASIC (ALP) kit). Analysis was performed using the cell density showing the least background in PBS injected mice.

### IAV micro-neutralisation assay

Micro-neutralisation (MN) assay was performed as described (Powell et al., 2012) with minor modifications. Briefly, a single cycle IAV expressing eGFP (S-eGFP (PR8)) containing the H1 haemagglutinin was titrated to give saturating infection of 3×10^4^ MDCK-SIAT1 cells per well in 96-well flat-bottom plates, detected by eGFP fluorescence. Murine sera were heat inactivated for 30 minutes at 56°C. Dilutions of sera were incubated with S-eGFP for 2 hours at 37°C before addition to 3×10^4^ MDCK-SIAT1 cells per well. Cells were then incubated overnight before fixing in 4% formaldehyde. The suppression of infection was measured on fixed cells by fluorescence on a CLARIOstar fluorescence plate reader.

### Vaccinia virus plaque assay

Ovaries collected in D0 (DMEM, 1% PenStrep) were homogenised using glass beads in screw cap tubes in a homogeniser (two cycles at speed 6.5 for 30 seconds). Samples were then placed on ice for 1-2 minutes and homogenisation was repeated. Samples were then subjected to three freeze-thaw cycles between 37°C and dry ice and sonicated three times for 30 seconds with 30 second intervals on ice. Supernatants containing virus were collected in new tubes after centrifugation at 10,000 rpm for 3 minutes at 4°C.

143B cells were seeded in 12 well plates at a density of 0.25×10^6^ cells per well in 1mL D10. The next day, log serial dilutions of virus-containing samples were prepared in D0. Supernatant was replaced with 550μl of diluted virus-containing samples and incubated for 2 hours at 37°C, swirling plates every 30 minutes to avoid drying. Virus containing samples were then removed and cells were covered in 1.5mL of D10 containing 1% Pen/Strep and 0.5% carboxymethylcellulose (CMC). 48 hours after infection, cells were carefully washed with PBS and fixed in 4% formaldehyde for 20 minutes at RT before staining with 0.5% crystal violet.

### Statistics

Statistical analysis was performed in GraphPad Prism v7.00 as detailed in the figure legends.

## Author contributions (using the CRediT taxonomy)

Conceptualisation: L.C., A.B., and J.R.; Methodology: L.C., A.B., T.K.T., J.F., I.P.-P. and T.P.; Software: n.a.; Validation: L.C. and J.R.; Formal analysis: L.C. and J.R.; Investigation: L.C., A.B., T.K.T. and J.F.; Resources: R.B. and P.B.; Data curation: L.C.; Writing – Original Draft: L.C. and J.R.; Writing – Review & Editing: all authors; Visualisation: L.C. and J.R.; Supervision: J.R., A.T., H.D. and P.B.; Project administration: L.C.; Funding acquisition: J.R.

## Acknowledgments

The authors thank Andrew McMichael, Adrian Hill, Daniel Radtke, Oliver Bannard, Nicolas Manel, Rachel Rigby and members of the Rehwinkel lab for discussion. The authors thank Uzi Gileadi and Vincenzo Cerundolo for their help with IAV infections. The authors thank Nicholas Proudfoot and Bernard Moss for providing respectively the 143B cells and the vVK1 vaccinia virus. The following reagents were obtained through the NIH AIDS Reagent Program, Division of AIDS, NIAID, NIH: HIV-1 Con B Gag Peptide Set, HIV-1 HXB2 Gag-EGFP Expression Vector (Cat#11468) from Dr. Marilyn Resh, HIV-1 NL4-3 ΔEnv EGFP Reporter Vector from Drs. Haili Zhang, Yan Zhou, and Robert Siliciano (cat# 11100), HIV-1IIIB pr55 Gag. This work was funded by the UK Medical Research Council [MRC core funding of the MRC Human Immunology Unit; J.R., J.F., H.D. and MRC Programme grant MR/K012037; P.B.], the Wellcome Trust [grant number 100954; J.R.], and the NIH, NIAID, DAIDS [UM1 grants AI00645 (Duke CHAVI-ID) and AI144371 (Duke CHAVD); P.B.]. P.B. is a Jenner Institute Investigator. Initial funding for the Virus Screening Facility was provided by the Oxford BRC and Cancer Research UK.The funders had no role in study design, data collection and analysis, decision to publish, or preparation of the manuscript.

## Declaration of interests

The authors have declared that no conflict of interest exits.

## Data availability statement

The authors declare that all data supporting the findings of this study are available within the paper and its supplementary information files.

## Figures and figure legends

**Fig S1:**
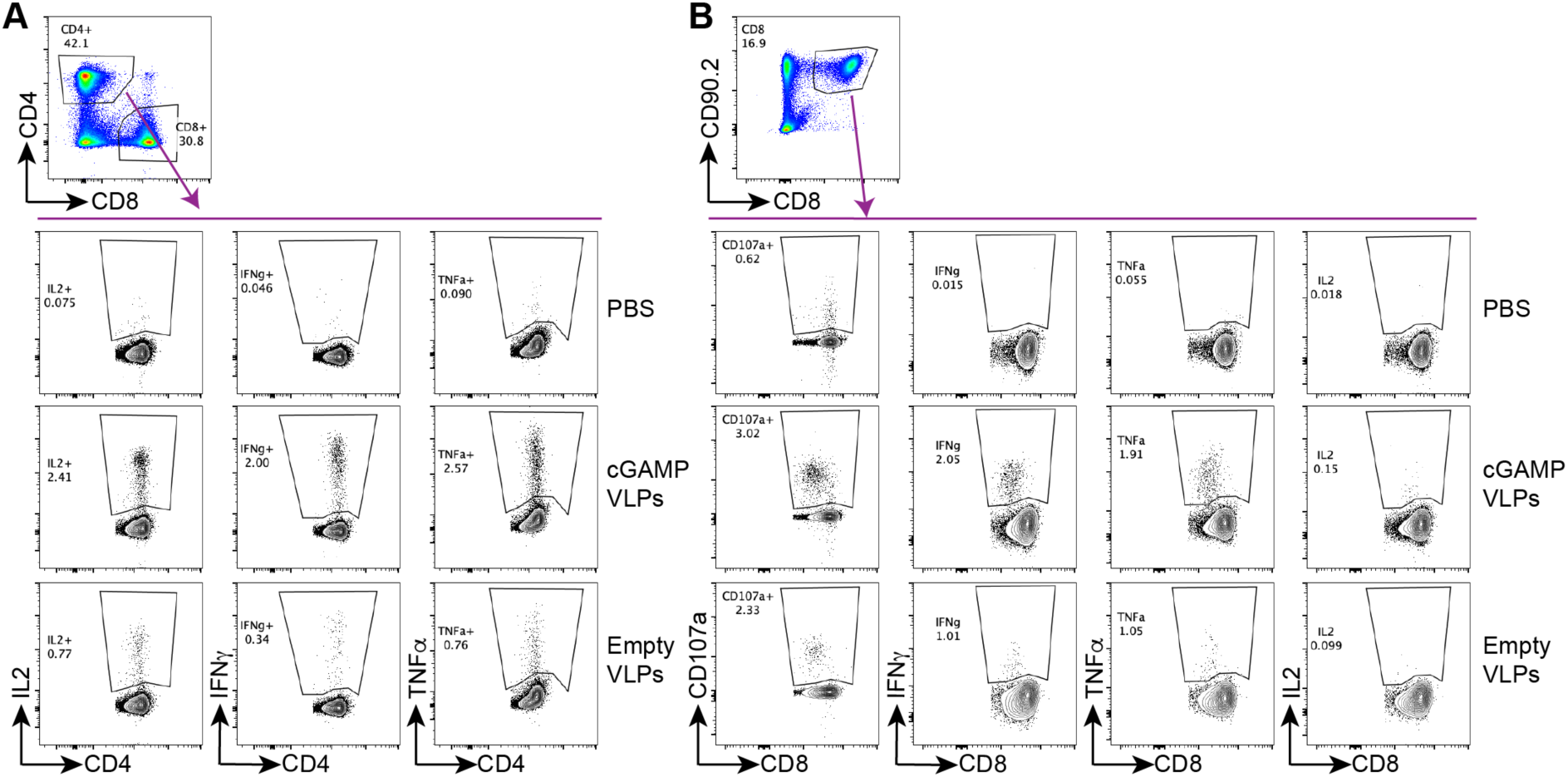
Gating strategies for the analysis of cytokine induction by Intracellular Cytokine Staining (ICS). C57BL/6 mice were injected with PBS as a control, cGAMP-VLPs or Empty-VLPs *via* the intra-muscular route. 14 days later, antigen-specific T cell responses were assessed by intracellular cytokine staining (ICS). **A.** For stimulation of CD4 T cells, BMMCs from C57BL/6 mice were pulsed overnight with cGAMP-VLPs and used to stimulate cells from spleens of immunised mice. Cells were co-cultured for six hours prior to evaluation of CD4 T cell responses by ICS. CD4 T cells were gated as live, MHC-II^-^, CD4^+^, CD8^-^. CD4 T cells expressing IL2, IFNγ or TNFα were analysed as shown. **B.** For stimulation of CD8 T cells, cells from spleens of immunised mice were stimulated with the HIV-SQV peptide for six hours prior to evaluation of CD8 T cell responses by ICS. CD8 T cells were gated as live, CD90.2^+^, CD8^+^. CD8 T cells expressing CD107a, IFNγ, TNFα or IL2 were analysed as shown.

**Fig S2:**
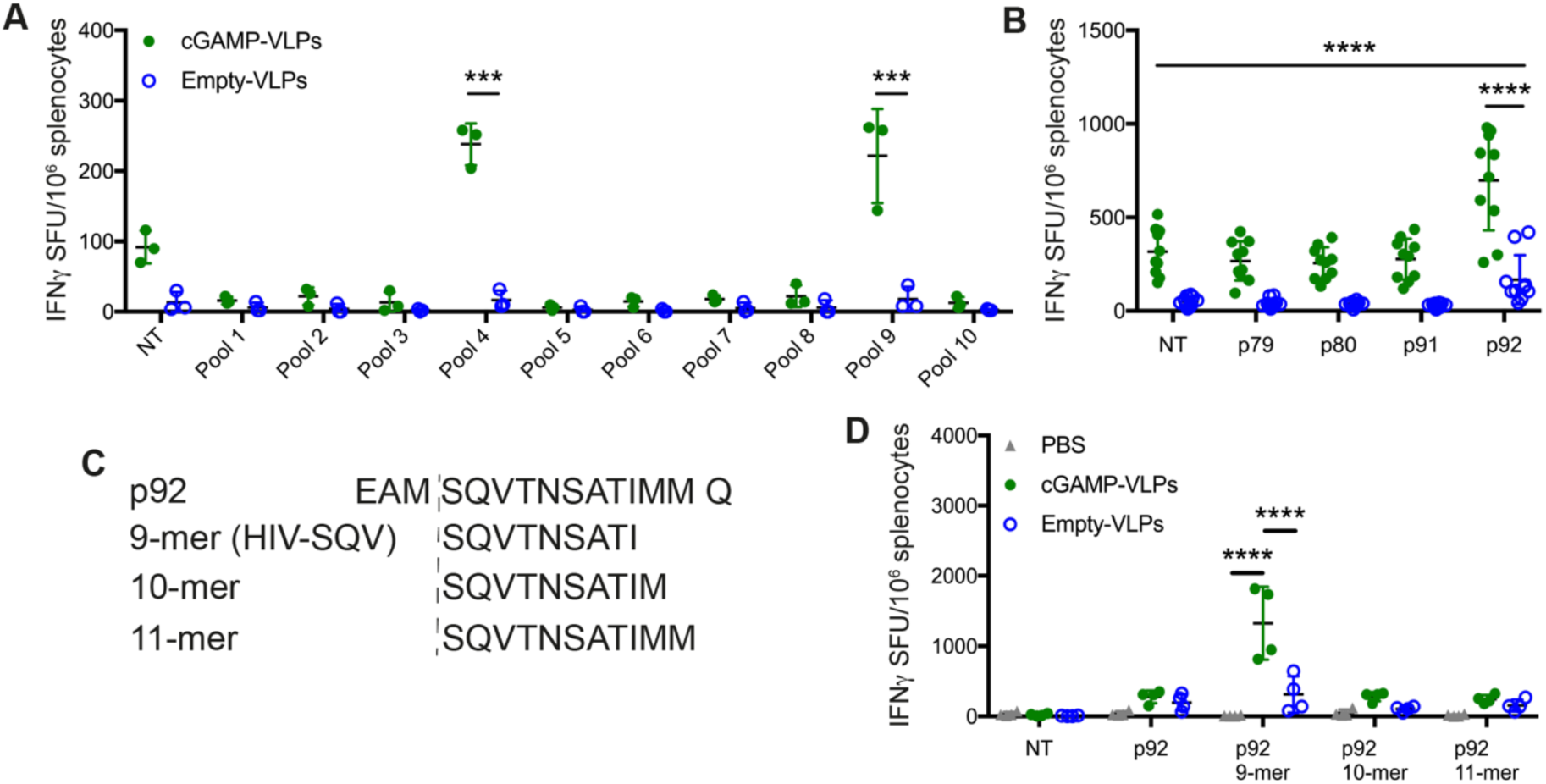
cGAMP-VLPs enhance T cell responses to the Gag HIV-SQV 9-mer peptide. C57BL/6 mice were injected with cGAMP-VLPs or Empty-VLPs *via* the intra-muscular route. 14 days later, antigen-specific T cell responses were assessed by IFNγ ELISPOT assay. **A.** Using a panel of 100 15-mer peptides spanning the HIV-1 Gag protein, we designed ten pools of 25 peptides so that each peptide was present in two pools and with minimal overlap between the pools. Cells from the spleens of immunised mice were stimulated for 24 hours with these peptide pools and responses were read out by IFNγ ELISPOT assay. **B.** The peptides that were common between pools 4 and 9 (p79, p80, p91, p92) were tested individually. **C.** Using NetMHC, we identified a 9-mer, a 10-mer and an 11-mer in p92 as predicted strong binders to H2-D^b^. **D.** Splenocytes from immunised mice were stimulated with the four versions of p92 shown in (C). Data in (A) and (D) are from a single experiment using three (A) or four (D) animals per group. Pooled data from two independent experiments including a total of 8 mice per group are shown in (B). In (A), (B) and (D), each symbol corresponds to one animal and mean and SD are shown. Statistical analyses were done using a 2-way ANOVA followed by Tukey’s multiple comparisons test, showing only selected comparisons. ***p<0.001; ****p<0.0001.

**Fig S3:**
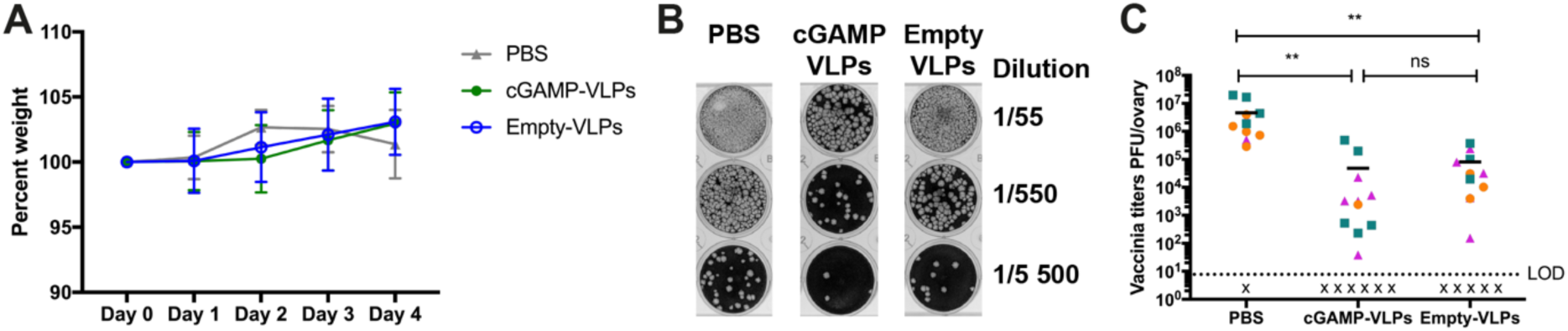
Evaluation of cGAMP-VLP-elicited protection in a vaccinia virus challenge model. Female C57BL/6 mice were injected with cGAMP-VLPs, Empty-VLPs, or PBS as a control *via* the intra-muscular route. One month later, mice were infected with 10^6^ PFU of a vaccinia virus expressing HIV Gag (vVK1) by intra-peritoneal inoculation. **A.** Weight loss was monitored over the course of infection and is shown as a percentage of weight prior to infection. **B-C.** Five days after infection, virus titres in the ovaries were quantified by plaque assay. A representative example of the plaque assay is shown in (B) and pooled data from three independent experiments including a total of 12-17 mice per group are shown in (C). A total of 12 mice (PBS) and 17 mice/group (cGAMP-VLPs and Empty-VLPs) were used in 3 independent experiments. In (A), mean and SD of pooled data are shown. In (C), each symbol represents data from an individual animal and colours indicate different experiments. Horizontal lines show the mean. x=sample below limit of detection (LOD). Statistical analyses were done using a 2-way ANOVA followed by Tukey’s multiple comparisons test. ns p≥0.05; **p<0.01.

**Fig S4:**
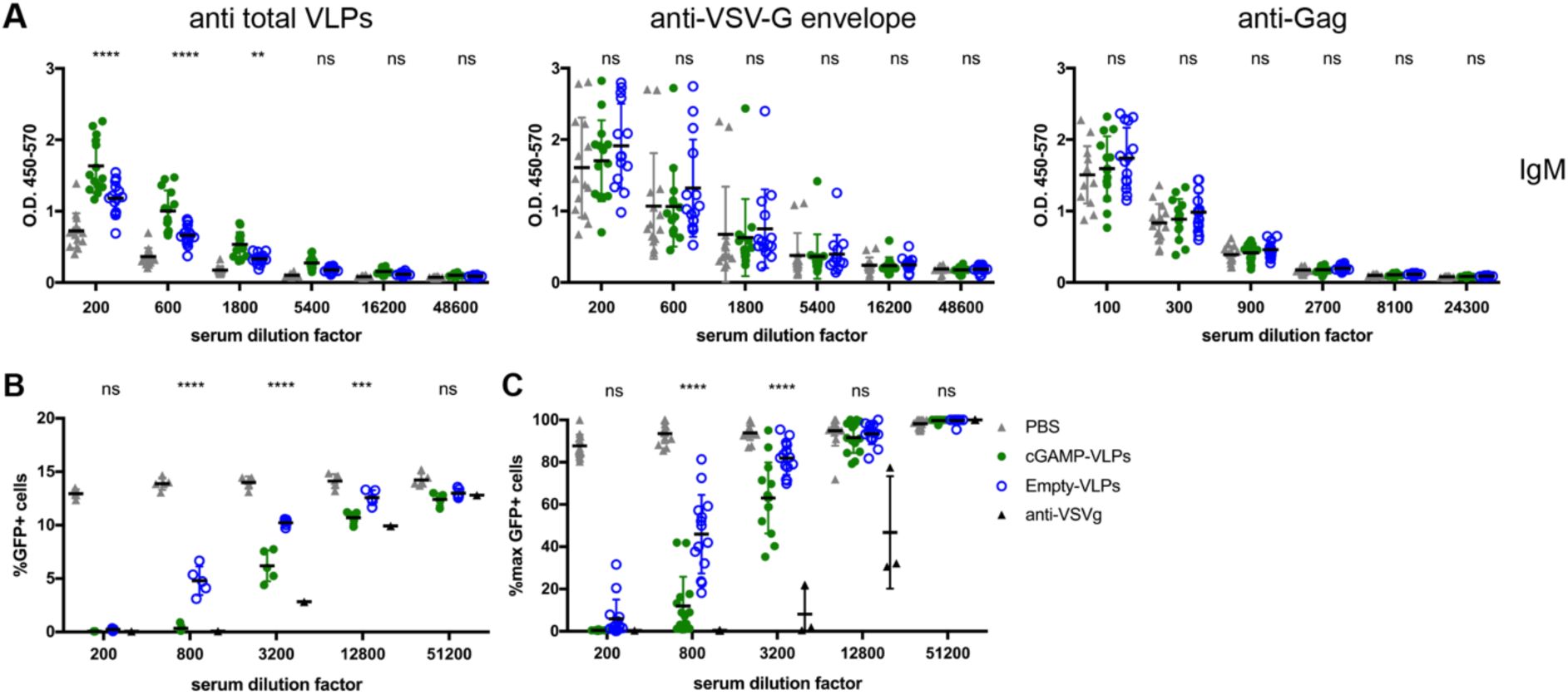
Immunisation with VLPs containing cGAMP increases neutralising antibody responses. C57BL/6 mice were injected with cGAMP-VLPs, Empty-VLPs, or PBS as a control *via* the intra-muscular route. 14 days later, serum antibody responses were evaluated. **A. IgM responses.** ELISA plates were coated with lysate from cGAMP-VLPs, recombinant VSV-G protein or recombinant HIV-1 Gag protein. IgM antibodies specific for these proteins were measured in sera from immunised mice. The optical density at increasing serum dilutions is shown. Data are pooled from three independent experiments. A total of 14 mice was analysed per condition. **B-C. cGAMP-VLPs enhance production of anti-VSV-G neutralising antibodies.** Serial dilutions of individual sera were incubated with VSV-G pseudotyped HIV-1-GFP for 90 minutes at 37°C before infection of HEK293 cells. As a control, dilutions of the anti-VSV-G neutralising antibody 8G5F11 were tested in parallel. After two days, infection was measured by quantifying GFP^+^ cells by flow cytometry. Data from a representative experiment is shown in (B). In (C), pooled data from three independent experiments including a total of 14 mice per condition are shown. For each experiment, the infection rate was normalised by setting the highest observed proportion of GFP^+^ cells to 100%. Symbols show data from individual animals, and the mean and SD are indicated. Statistical analyses were done using a 2-way ANOVA followed by Tukey’s multiple comparisons test, only showing significance between cGAMP-VLPs and Empty-VLPs. ns p≥0.05; *p<0.05; **p<0.01; ***p<0.001; ****p<0.0001.

**Fig S5:**
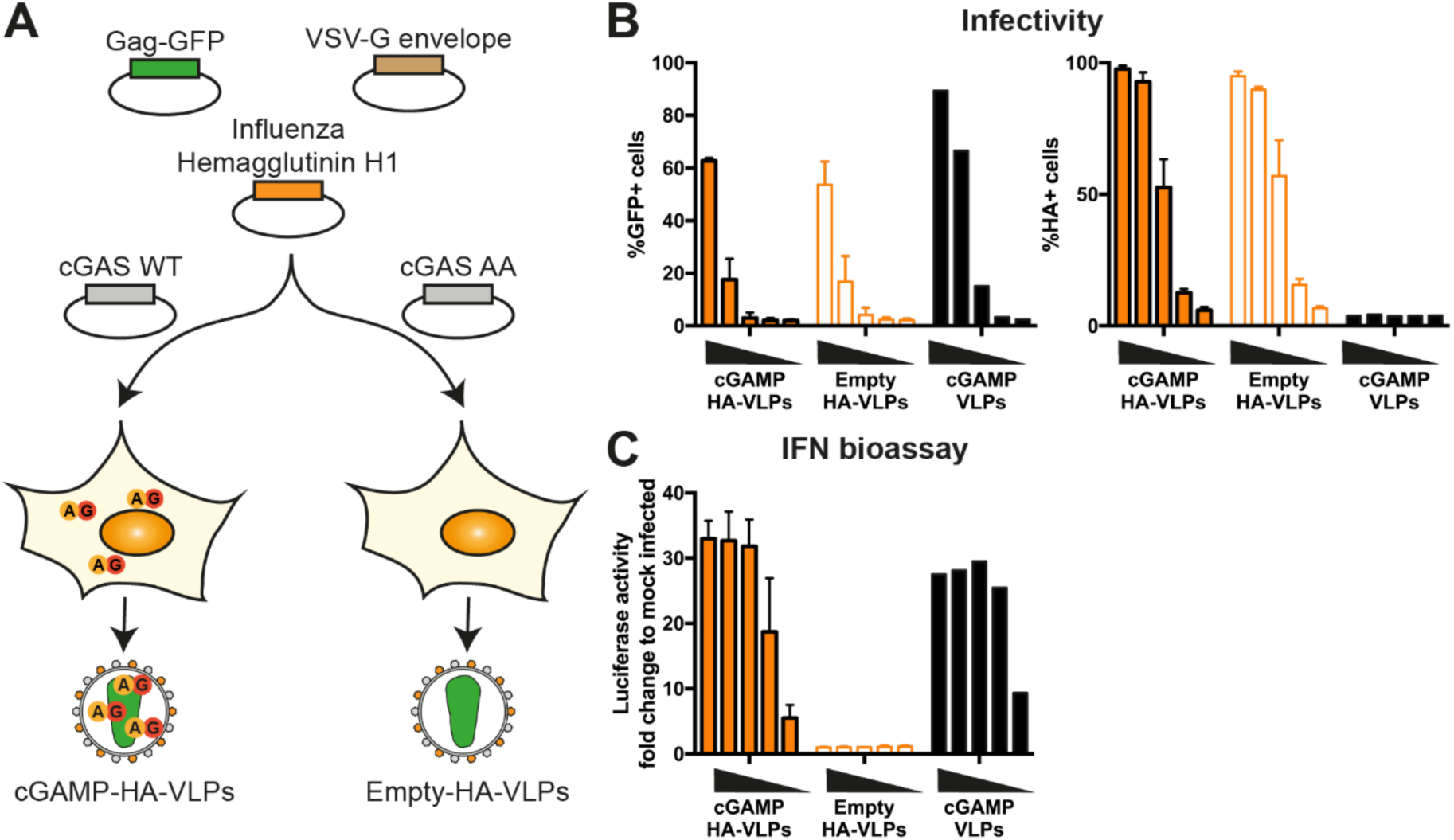
Pseudotyping of cGAMP-VLPs with IAV Haemagglutinin (HA). **A. Schematic representation of cGAMP-HA-VLP and Empty-HA-VLP production.** HEK293T cells were transfected with plasmids encoding HIV-1 Gag-GFP, VSV-G envelope and IAV HA. cGAMP-HA-VLPs were collected from cells co-expressing cGAS WT and Empty-HA-VLPs from cells co-expressing catalytically inactive cGAS AA. **B. IAV HA is present in HA-VLPs.** HEK293 cells were infected with decreasing amounts of cGAMP-HA-VLPs and Empty-HA-VLPs (1/5 serial dilutions starting at 2μL of VLP stocks per well). Infection was monitored 24 hours later by quantifying GFP^+^ and HA^+^ cells by flow cytometry. cGAMP-VLPs were used for comparison. **C. cGAMP-HA-VLPs induce a similar IFN-I response in infected cells compared to cGAMP-VLPs.** Supernatants from infected cells shown in (B) were tested for the presence of IFN-I as shown in Fig 1C. Data in (B) and (C) are pooled from two independent HA-VLP productions tested simultaneously in technical duplicates in infectivity and IFN-I bioassays; mean and SD are shown.

**Supplementary Table 1.**
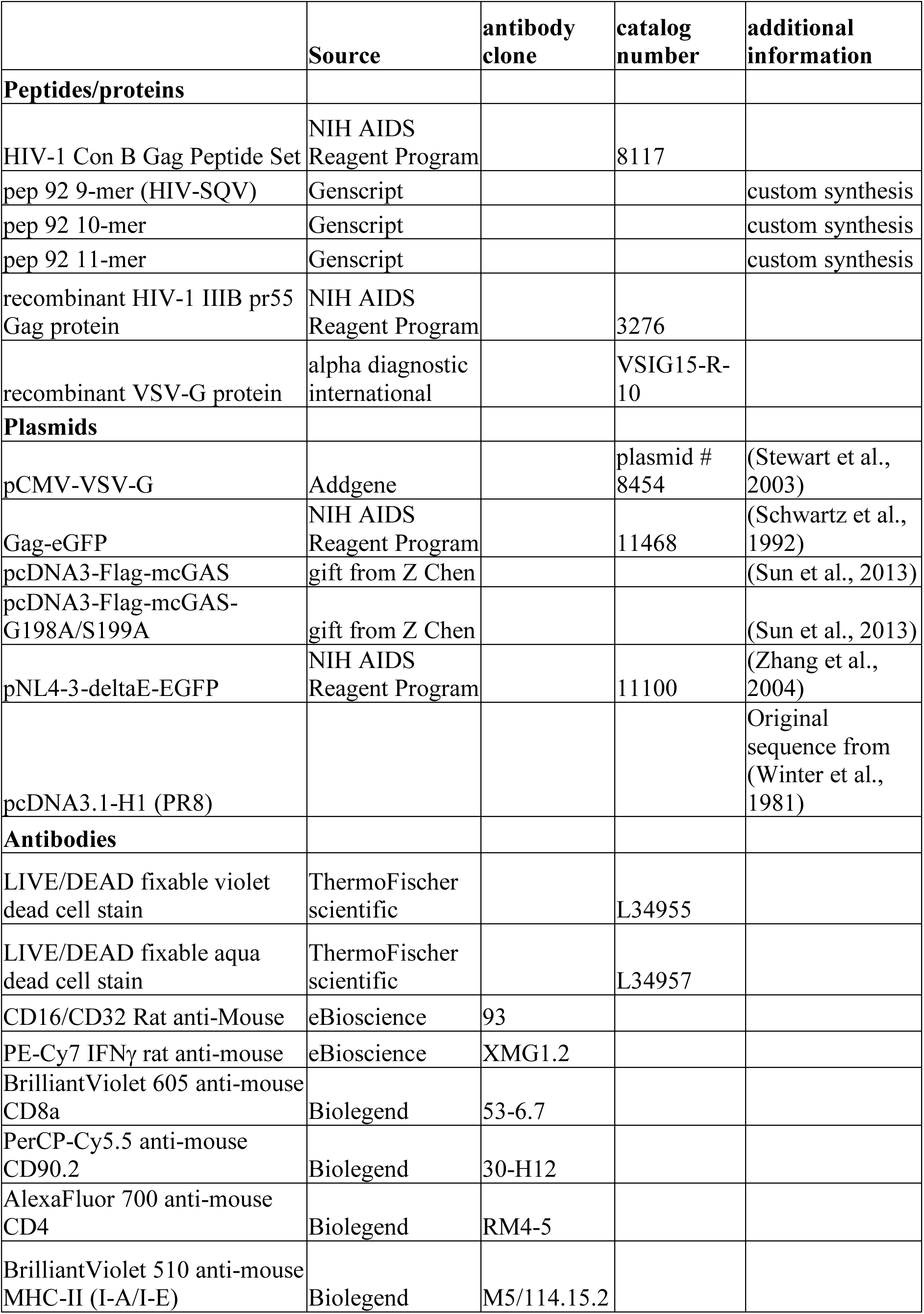
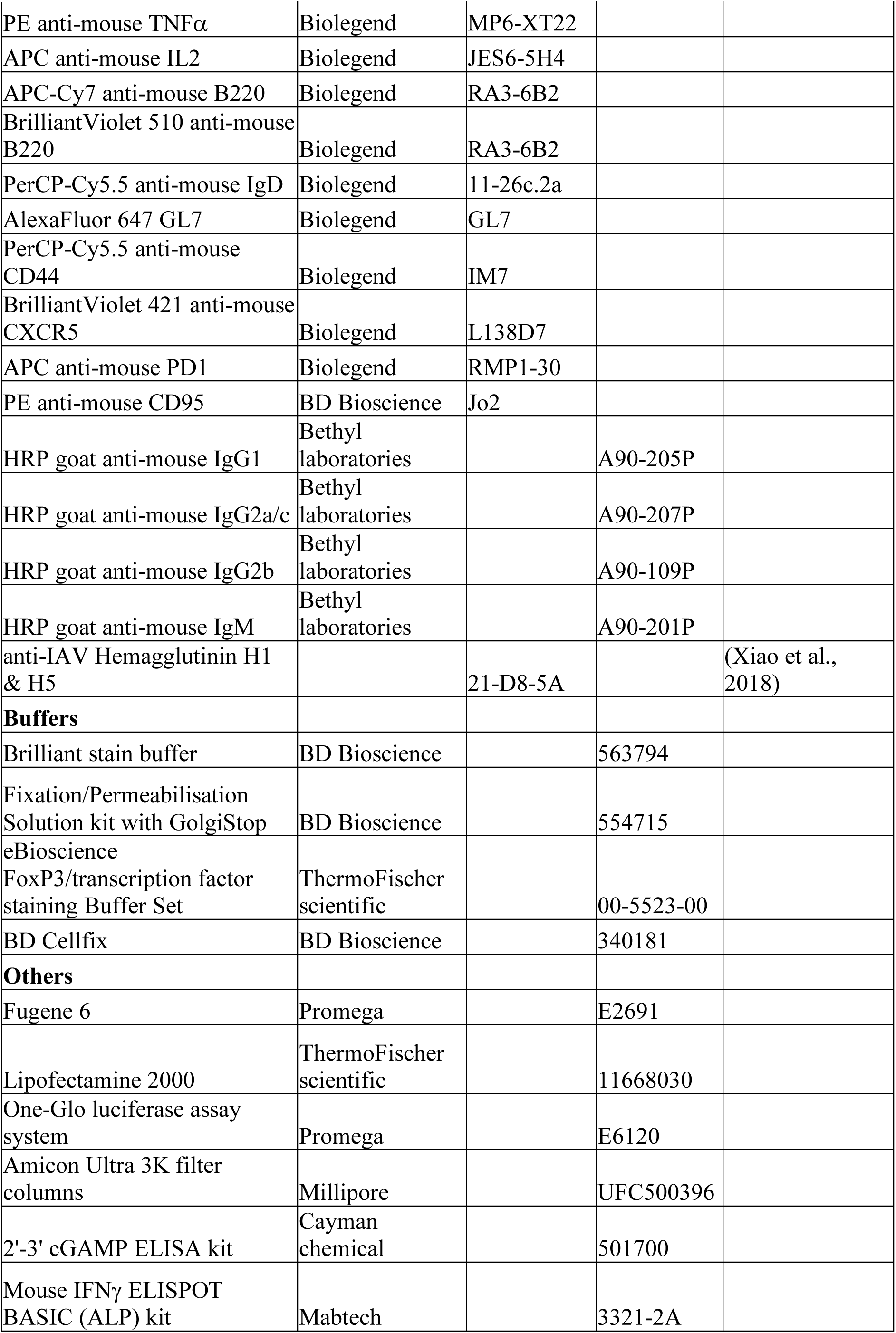
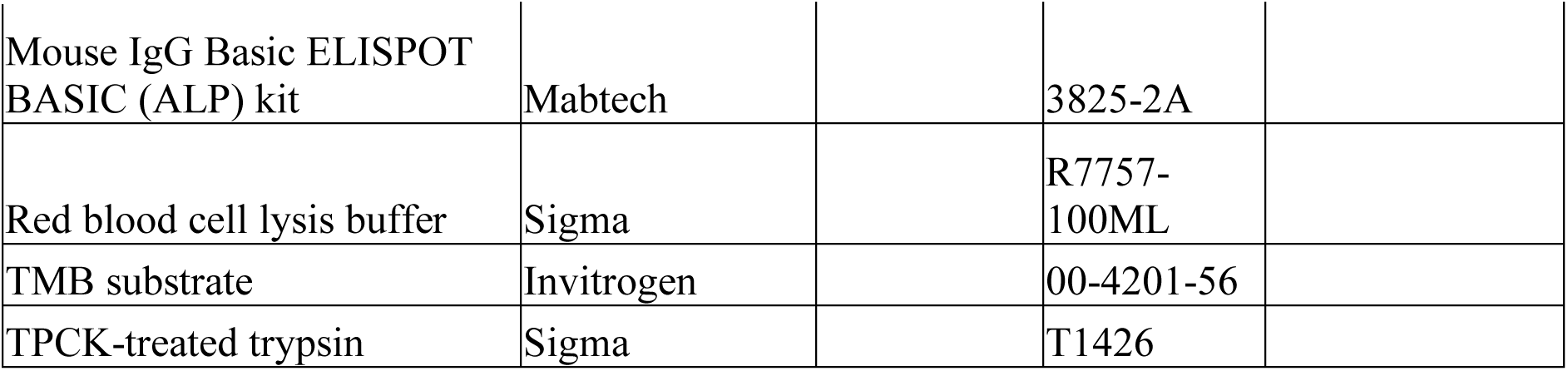
Reagents

## References

Ablasser, A., M. Goldeck, T. Cavlar, T. Deimling, G. Witte, I. Rohl, K.P. Hopfner, J. Ludwig, and V. Hornung. 2013. cGAS produces a 2’-5’-linked cyclic dinucleotide second messenger that activates STING. Nature 498:380–384.

Blaauboer, S.M., V.D. Gabrielle, and L. Jin. 2014. MPYS/STING-mediated TNF-alpha, not type I IFN, is essential for the mucosal adjuvant activity of (3’-5’)-cyclic-di-guanosine-monophosphate in vivo. J Immunol 192:492–502.

Blaauboer, S.M., S. Mansouri, H.R. Tucker, H.L. Wang, V.D. Gabrielle, and L. Jin. 2015. The mucosal adjuvant cyclic di-GMP enhances antigen uptake and selectively activates pinocytosis-efficient cells in vivo. Elife 4:e06670.

Borrow, P., H. Lewicki, B.H. Hahn, G.M. Shaw, and M.B. Oldstone. 1994. Virus-specific CD8+ cytotoxic T-lymphocyte activity associated with control of viremia in primary human immunodeficiency virus type 1 infection. J Virol 68:6103–6110.

Bridgeman, A., J. Maelfait, T. Davenne, T. Partridge, Y. Peng, A. Mayer, T. Dong, V. Kaever, P. Borrow, and J. Rehwinkel. 2015. Viruses transfer the antiviral second messenger cGAMP between cells. Science 349:1228–1232.

Burdette, D.L., K.M. Monroe, K. Sotelo-Troha, J.S. Iwig, B. Eckert, M. Hyodo, Y. Hayakawa, and R.E. Vance. 2011. STING is a direct innate immune sensor of cyclic di-GMP. Nature 478:515–518.

Cai, X., Y.H. Chiu, and Z.J. Chen. 2014. The cGAS-cGAMP-STING pathway of cytosolic DNA sensing and signaling. Mol Cell 54:289–296.

Carozza, J.A., V. Böhnert, K.C. Nguyen, G. Skariah, K.E. Shaw, J.A. Brown, M. Rafat, R. von Eyben, E.E. Graves, J.S. Glenn, M. Smith, and L. Li. 2019. Extracellular 2’3’-cGAMP is an immunotransmitter produced by cancer cells and regulated by ENPP1. bioRxiv 539312.

Coffman, R.L., A. Sher, and R.A. Seder. 2010. Vaccine adjuvants: putting innate immunity to work. Immunity 33:492–503.

Corrales, L., L.H. Glickman, S.M. McWhirter, D.B. Kanne, K.E. Sivick, G.E. Katibah, S.R. Woo, E. Lemmens, T. Banda, J.J. Leong, K. Metchette, T.W. Dubensky, Jr., and T.F. Gajewski. 2015. Direct Activation of STING in the Tumor Microenvironment Leads to Potent and Systemic Tumor Regression and Immunity. Cell Rep 11:1018–1030.

Crotty, S. 2019. T Follicular Helper Cell Biology: A Decade of Discovery and Diseases. Immunity 50:1132–1148.

Cucak, H., U. Yrlid, B. Reizis, U. Kalinke, and B. Johansson-Lindbom. 2009. Type I interferon signaling in dendritic cells stimulates the development of lymph-node-resident T follicular helper cells. Immunity 31:491–501.

Cyster, J.G., and C.D.C. Allen. 2019. B Cell Responses: Cell Interaction Dynamics and Decisions. Cell 177:524–540.

Demaria, O., A. De Gassart, S. Coso, N. Gestermann, J. Di Domizio, L. Flatz, O. Gaide, O. Michielin, P. Hwu, T.V. Petrova, F. Martinon, R.L. Modlin, D.E. Speiser, and M. Gilliet. 2015. STING activation of tumor endothelial cells initiates spontaneous and therapeutic antitumor immunity. Proc Natl Acad Sci U S A 112:15408–15413.

Deml, L., C. Speth, M.P. Dierich, H. Wolf, and R. Wagner. 2005. Recombinant HIV-1 Pr55gag virus-like particles: potent stimulators of innate and acquired immune responses. Mol Immunol 42:259–277.

Diner, E.J., D.L. Burdette, S.C. Wilson, K.M. Monroe, C.A. Kellenberger, M. Hyodo, Y. Hayakawa, M.C. Hammond, and R.E. Vance. 2013. The innate immune DNA sensor cGAS produces a noncanonical cyclic dinucleotide that activates human STING. Cell Rep 3:1355–1361.

Dubensky, T.W., Jr., D.B. Kanne, and M.L. Leong. 2013. Rationale, progress and development of vaccines utilizing STING-activating cyclic dinucleotide adjuvants. Ther Adv Vaccines 1:131–143.

Ebensen, T., R. Libanova, K. Schulze, T. Yevsa, M. Morr, and C.A. Guzman. 2011. Bis-(3’,5’)-cyclic dimeric adenosine monophosphate: strong Th1/Th2/Th17 promoting mucosal adjuvant. Vaccine 29:5210–5220.

Gentili, M., J. Kowal, M. Tkach, T. Satoh, X. Lahaye, C. Conrad, M. Boyron, B. Lombard, S. Durand, G. Kroemer, D. Loew, M. Dalod, C. Thery, and N. Manel. 2015. Transmission of innate immune signaling by packaging of cGAMP in viral particles. Science 349:1232–1236.

Hansen, S.G., J.C. Ford, M.S. Lewis, A.B. Ventura, C.M. Hughes, L. Coyne-Johnson, N. Whizin, K. Oswald, R. Shoemaker, T. Swanson, A.W. Legasse, M.J. Chiuchiolo, C.L. Parks, M.K. Axthelm, J.A. Nelson, M.A. Jarvis, M. Piatak, Jr., J.D. Lifson, and L.J. Picker. 2011. Profound early control of highly pathogenic SIV by an effector memory T-cell vaccine. Nature 473:523–527.

Holechek, S.A., M.S. McAfee, L.M. Nieves, V.P. Guzman, K. Manhas, T. Fouts, K. Bagley, and J.N. Blattman. 2016. Retinaldehyde dehydrogenase 2 as a molecular adjuvant for enhancement of mucosal immunity during DNA vaccination. Vaccine 34:5629–5635.

Hong, S., Z. Zhang, H. Liu, M. Tian, X. Zhu, Z. Zhang, W. Wang, X. Zhou, F. Zhang, Q. Ge, B. Zhu, H. Tang, Z. Hua, and B. Hou. 2018. B Cells Are the Dominant Antigen-Presenting Cells that Activate Naive CD4(+) T Cells upon Immunization with a Virus-Derived Nanoparticle Antigen. Immunity 49:695–708 e694.

Itano, A.A., and M.K. Jenkins. 2003. Antigen presentation to naive CD4 T cells in the lymph node. Nat Immunol 4:733–739.

Joffre, O., M.A. Nolte, R. Sporri, and C. Reis e Sousa. 2009. Inflammatory signals in dendritic cell activation and the induction of adaptive immunity. Immunol Rev 227:234–247.

Karacostas, V., K. Nagashima, M.A. Gonda, and B. Moss. 1989. Human immunodeficiency virus-like particles produced by a vaccinia virus expression vector. Proc Natl Acad Sci U S A 86:8964–8967.

Krammer, F. 2019. The human antibody response to influenza A virus infection and vaccination. Nat Rev Immunol 19:383–397.

Kuse, N., X. Sun, T. Akahoshi, A. Lissina, T. Yamamoto, V. Appay, and M. Takiguchi. 2019. Priming of HIV-1-specific CD8(+) T cells with strong functional properties from naive T cells. EBioMedicine 42:109–119.

Li, L., Q. Yin, P. Kuss, Z. Maliga, J.L. Millan, H. Wu, and T.J. Mitchison. 2014. Hydrolysis of 2’3’-cGAMP by ENPP1 and design of nonhydrolyzable analogs. Nat Chem Biol 10:1043–1048.

Li, T., H. Cheng, H. Yuan, Q. Xu, C. Shu, Y. Zhang, P. Xu, J. Tan, Y. Rui, P. Li, and X. Tan. 2016. Antitumor Activity of cGAMP via Stimulation of cGAS-cGAMP-STING-IRF3 Mediated Innate Immune Response. Sci Rep 6:19049.

Li, X.D., J. Wu, D. Gao, H. Wang, L. Sun, and Z.J. Chen. 2013. Pivotal roles of cGAS-cGAMP signaling in antiviral defense and immune adjuvant effects. Science 341:1390–1394.

Linterman, M.A., and D.L. Hill. 2016. Can follicular helper T cells be targeted to improve vaccine efficacy? F1000Res 5:

Mayer, A., J. Maelfait, A. Bridgeman, and J. Rehwinkel. 2017. Purification of Cyclic GMP-AMP from Viruses and Measurement of Its Activity in Cell Culture. Methods Mol Biol 1656:143–152.

Milone, M.C., and U. O’Doherty. 2018. Clinical use of lentiviral vectors. Leukemia 32:1529–1541.

Nielsen, M., and M. Andreatta. 2016. NetMHCpan-3.0; improved prediction of binding to MHC class I molecules integrating information from multiple receptor and peptide length datasets. Genome Med 8:33.

Nurieva, R.I., Y. Chung, G.J. Martinez, X.O. Yang, S. Tanaka, T.D. Matskevitch, Y.H. Wang, and C. Dong. 2009. Bcl6 mediates the development of T follicular helper cells. Science 325:1001–1005.

O’Garra, A. 1998. Cytokines induce the development of functionally heterogeneous T helper cell subsets. Immunity 8:275–283.

Panagioti, E., P. Klenerman, L.N. Lee, S.H. van der Burg, and R. Arens. 2018. Features of Effective T Cell-Inducing Vaccines against Chronic Viral Infections. Front Immunol 9:276.

Pelegrin, M., M. Naranjo-Gomez, and M. Piechaczyk. 2015. Antiviral Monoclonal Antibodies: Can They Be More Than Simple Neutralizing Agents? Trends Microbiol 23:653–665.

Powell, T.J., J.D. Silk, J. Sharps, E. Fodor, and A.R. Townsend. 2012. Pseudotyped influenza A virus as a vaccine for the induction of heterotypic immunity. J Virol 86:13397–13406.

Rappuoli, R., C.W. Mandl, S. Black, and E. De Gregorio. 2011. Vaccines for the twenty-first century society. Nat Rev Immunol 11:865–872.

Reed, L.J., and H. Muench. 1938. A simple method of estimating fifty per cent endpoints12. American Journal of Epidemiology 27:493–497.

Riteau, N., A.J. Radtke, K. Shenderov, L. Mittereder, S.D. Oland, S. Hieny, D. Jankovic, and A. Sher. 2016. Water-in-Oil-Only Adjuvants Selectively Promote T Follicular Helper Cell Polarization through a Type I IFN and IL-6-Dependent Pathway. J Immunol 197:3884–3893.

Sage, P.T., L.M. Francisco, C.V. Carman, and A.H. Sharpe. 2013. The receptor PD-1 controls follicular regulatory T cells in the lymph nodes and blood. Nat Immunol 14:152–161.

Schwartz, S., M. Campbell, G. Nasioulas, J. Harrison, B.K. Felber, and G.N. Pavlakis. 1992. Mutational inactivation of an inhibitory sequence in human immunodeficiency virus type 1 results in Rev-independent gag expression. J Virol 66:7176–7182.

Shi, S., H. Zhu, X. Xia, Z. Liang, X. Ma, and B. Sun. 2019. Vaccine adjuvants: Understanding the structure and mechanism of adjuvanticity. Vaccine 37:3167–3178.

Stewart, S.A., D.M. Dykxhoorn, D. Palliser, H. Mizuno, E.Y. Yu, D.S. An, D.M. Sabatini, I.S. Chen, W.C. Hahn, P.A. Sharp, R.A. Weinberg, and C.D. Novina. 2003. Lentivirus-delivered stable gene silencing by RNAi in primary cells. RNA 9:493–501.

Sun, L., J. Wu, F. Du, X. Chen, and Z.J. Chen. 2013. Cyclic GMP-AMP synthase is a cytosolic DNA sensor that activates the type I interferon pathway. Science 339:786–791.

Temizoz, B., E. Kuroda, and K.J. Ishii. 2018. Combination and inducible adjuvants targeting nucleic acid sensors. Curr Opin Pharmacol 41:104–113.

Trumpfheller, C., J.S. Finke, C.B. Lopez, T.M. Moran, B. Moltedo, H. Soares, Y. Huang, S.J. Schlesinger, C.G. Park, M.C. Nussenzweig, A. Granelli-Piperno, and R.M. Steinman. 2006. Intensified and protective CD4+ T cell immunity in mice with anti-dendritic cell HIV gag fusion antibody vaccine. J Exp Med 203:607–617.

Wang, H., S. Hu, X. Chen, H. Shi, C. Chen, L. Sun, and Z.J. Chen. 2017. cGAS is essential for the antitumor effect of immune checkpoint blockade. Proc Natl Acad Sci U S A 114:1637–1642.

Wang, J., P. Li, and M.X. Wu. 2016. Natural STING Agonist as an “Ideal” Adjuvant for Cutaneous Vaccination. J Invest Dermatol 136:2183–2191.

Winter, G., S. Fields, and G.G. Brownlee. 1981. Nucleotide sequence of the haemagglutinin gene of a human influenza virus H1 subtype. Nature 292:72–75.

Xiao, J.H., P. Rijal, L. Schimanski, A.K. Tharkeshwar, E. Wright, W. Annaert, and A. Townsend. 2018. Characterization of Influenza Virus Pseudotyped with Ebolavirus Glycoprotein. J Virol 92:

Xu, R., A.J. Johnson, D. Liggitt, and M.J. Bevan. 2004. Cellular and humoral immunity against vaccinia virus infection of mice. J Immunol 172:6265–6271.

Yang, L., H. Yang, K. Rideout, T. Cho, K.I. Joo, L. Ziegler, A. Elliot, A. Walls, D. Yu, D. Baltimore, and P. Wang. 2008. Engineered lentivector targeting of dendritic cells for in vivo immunization. Nat Biotechnol 26:326–334.

Zhang, H., Y. Zhou, C. Alcock, T. Kiefer, D. Monie, J. Siliciano, Q. Li, P. Pham, J. Cofrancesco, D. Persaud, and R.F. Siliciano. 2004. Novel single-cell-level phenotypic assay for residual drug susceptibility and reduced replication capacity of drug-resistant human immunodeficiency virus type 1. J Virol 78:1718–1729.

